# Alternative splicing of its 5’-UTR limits CD20 mRNA translation and enables resistance to CD20-directed immunotherapies

**DOI:** 10.1101/2023.02.19.529123

**Authors:** Zhiwei Ang, Luca Paruzzo, Katharina E. Hayer, Carolin Schmidt, Manuel Torres Diz, Feng Xu, Urvi Zankharia, Yunlin Zhang, Samantha Soldan, Sisi Zheng, Catherine D. Falkenstein, Joseph P. Loftus, Scarlett Y. Yang, Mukta Asnani, Patricia King Sainos, Vinodh Pillai, Emeline Chong, Marilyn M. Li, Sarah K. Tasian, Yoseph Barash, Paul M. Lieberman, Marco Ruella, Stephen J. Schuster, Andrei Thomas-Tikhonenko

## Abstract

Aberrant skipping of coding exons in CD19 and CD22 compromises responses to immunotherapy for B-cell malignancies. Here, we show that the *MS4A1* gene encoding human CD20 also produces several mRNA isoforms with distinct 5’ untranslated regions (5’-UTR). Four variants (V1-4) were detectable by RNA-seq in distinct stages of normal B-cell differentiation and B-lymphoid malignancies, with V1 and V3 being the most abundant by far. During B-cell activation and Epstein-Barr virus infection, redirection of splicing from V1 to V3 coincided with increased CD20 positivity. Similarly, in diffuse large B-cell lymphoma only V3, but not V1, correlated with CD20 protein levels, suggesting that V1 might be translation-deficient. Indeed, the longer V1 isoform was found to contain upstream open reading frames (uORFs) and a stem-loop structure, which cooperatively inhibited polysome recruitment. By modulating CD20 isoforms with splice-switching Morpholino oligomers, we enhanced CD20 expression and anti-CD20 antibody rituximab-mediated cytotoxicity in a panel of B-cell lines. Furthermore, reconstitution of CD20-knockout cells with V3 mRNA led to the recovery of CD20 positivity, while V1-reconstituted cells had undetectable levels of CD20 protein. Surprisingly, *in vitro* CD20-directed CAR T cells were able to kill both V3- and V1-expressing cells, but the bispecific T cell engager mosunetuzumab was only effective against V3-expressing cells. To determine whether CD20 splicing is involved in immunotherapy resistance, we performed RNA-seq on four post-mosunetuzumab follicular lymphoma relapses and discovered that in two of them downregulation of CD20 was accompanied by the V3-to-V1 shift. Thus, splicing-mediated mechanisms of epitope loss extend to CD20-directed immunotherapies.

**Key Points:**

1. In normal & malignant human B cells, CD20 mRNA is alternatively spliced into four 5’-UTR isoforms, some of which are translation-deficient.
2. The balance between translation-deficient and -competent isoforms modulates CD20 protein levels & responses to CD20-directed immunotherapies

**Explanation of Novelty:** We discovered that in normal and malignant B-cells, CD20 mRNA is alternatively spliced to generate four distinct 5’-UTRs, including the longer translation-deficient V1 variant. Cells predominantly expressing V1 were still sensitive to CD20-targeting chimeric antigen receptor T-cells. However, they were resistant to the bispecific anti-CD3/CD20 antibody mosunetuzumab, and the shift to V1 were observed in CD20-negative post-mosunetuzumab relapses of follicular lymphoma.

## INTRODUCTION

The plasma membrane protein CD20, encoded in humans by the *MS4A1* gene, is a clinically significant target for multiple monoclonal antibody (mAb) therapies, due to its lineage-specific expression on mature B-cells and the malignancies derived therefrom ^1^. In adult patients, CD20-directed therapies are administered alone or in combination with chemotherapy as standards of care for mature B-cell neoplasms such as diffuse large B-cell lymphoma (DLBCL), follicular lymphoma (FL), Burkitt lymphoma (BL), and high-grade B-cell lymphomas (HGBCL) ^2,3^. By far the most widely prescribed anti-CD20 mAb is rituximab, which is on the World Health Organization’s List of Essential Medicines ^4^. In pediatric patients, rituximab and chemotherapy combinations are approved for previously untreated, advanced stage CD20-positive DLBCL, BL, and HGBCL ^5^. In EU and other countries, rituximab is also used to treat children and adults with CD20-positive B-cell lymphoblastic leukemia (B-ALL) ^6^. Newer CD20-directed immunotherapies include mosunetuzumab, a CD20xCD3 bispecific monoclonal antibody that redirects T cells to engage and eliminate malignant B-cells ^7,8^. In 2022, mosunetuzumab was granted accelerated approval in the EU and by the FDA for the treatment of relapsed or refractory (r/r) FL as a third-line or later therapy ^9^.

While CD20-targeted immunotherapies have increased the median overall survival of patients with B-cell malignancies, de novo or acquired immunotherapy resistance due to antigen loss remains a challenge ^2,10^. In patients with r/r B-cell NHLs (B-NHL), low levels of CD20 at baseline (16/293 or 5.5% of cases) was associated with a lack of response to mosunetuzumab monotherapy. In chronic lymphocytic leukemia (CLL), CD20 levels are generally even lower relative to other mature B-cell neoplasms ^11-14^. This is believed to contribute to the relatively low efficacy of rituximab monotherapy against CLL ^15^. Beyond this intrinsic resistance, CD20 protein was lost in 7/26 (27%) of patients with B-NHL that relapsed after initial responses to mosunetuzumab ^16^. Overall, up to a third of all adult patients with DLBCL, and the majority of patients with FL and (CLL) are not cured by rituximab +/-chemotherapy combinations ^17,18^. This resistance is commonly associated with the down-modulation of CD20 ^19-24^.

Despite the clear role of CD20 antigen loss in immunotherapy resistance, significant gaps in our understanding of the underlying mechanism(s) still exist. A recent report demonstrated that in a Burkitt lymphoma cell line, acute loss of CD20 reduces CD19 expression over time ^25^, although this was not an immediate effect, and CD19 loss in CD20-negative relapses has not been reported. We previously reported that the Myc oncoprotein down-regulates CD20 mRNA expression in human B-cells and renders them partly resistant to rituximab *in vitro* ^26^. What, if any, role this regulation plays in patients with B-cell lymphomas remains to be determined. The loss of CD20 protein in 3/4 of mosunetuzumab-resistant B-NHL tumors analyzed using next-generation sequencing could not be explained by the disappearance of CD20 mRNA or the emergence of *MS4A1* genetic variants, as recently reported ^16^. A lack of concordance between CD20 mRNA and protein was also observed in CLL samples, which were found to be low in CD20 protein despite a near normal level of CD20 mRNA relative to normal B-cells ^27^. Here we addressed posttranscriptional mechanisms of CD20 dysregulation, with focus on alternative splicing.

## MATERIALS AND METHODS

### Dataset usage

The eBL2 (PRJNA292327), eBL3(PRJNA374464) [Ref ^28^], DLBCL2(PRJNA531552), DLBCL3 (PRJNA752102), DLBCL4 (PRJNA373954) [Ref ^29^], FL1(PRJNA596663) [Ref ^30^], FL2(PRJNA263567) [Ref ^31^], CLL2(PRJNA376727) [Ref ^32^], CLL3(PRJEB4498), CLL4(PRJNA792609) [Ref ^33^], CLL5(PRJNA450999) [Ref ^34^], bone marrow subsets (PRJNA475684) [Ref ^35-37^] and peripheral blood B cell (PRJNA418779) [Ref ^38^] datasets were from the BioProject databases of the National Center for Biotechnology Information (NCBI). The eBL1(phs001282.V2.p1) and CLL1(phs000767.V1.p1) [Ref ^39^] datasets were from the dbGaP database of the NCBI. For the TCGA DLBCL samples, CD20 isoform RNA level RSEM data and protein RPPA data were obtained from the TSVdb (http://www.tsvdb.com/) ^40^ and cBioPortal (https://www.cbioportal.org/) ^41^ websites respectively. Controlled access datasets were downloaded under the dbGaP Project #11199: “Posttranscriptional regulation in B-lymphoid malignancies”.

### RNA-seq analysis

RNA-seq reads were first trimmed to remove adapters (BBTools v38.96) and were then aligned using STAR v2.7.9a to the hg38 reference genome while providing known gene isoforms through the GENCODE annotation V32. In addition, we used STAR flags “–quantMode GeneCounts” and “– alignSJoverhangMin 8” to quantify genes and ensure spliced reads had an overhang of at least eight bases. Junction spanning reads were obtained from the STAR “*_SJ.out.tab” result files, and each entry was normalized by dividing by the total number of junction-spanning reads and multiplying by a factor of 1 million to obtain the “junctions per million” or JPM. Visualization and downstream analyses were conducted in R using the ggplot2 and tidyverse packages. TPMs for all samples were calculated using TPMCalculator ^42^ version 0.0.3 followed by ComBat-seq ^43^ batch correction as implemented in the R Bioconductor package sva (version 3.42.0).

### MAJIQ analysis

For RNA splicing quantification, Majiq v2.4.dev+g85d07819 was used. The bam files from STAR and the gencode.v37 annotation file were processed with the MAJIQ-build functionality to generate the splicegraph and MAJIQ files. Next, for each patient sample, the files from the corresponding pre- and post-relapse samples were compared with the MAJIQ deltapsi tool to estimate changes in splicing (deltaPSI). Results were exported with voila tsv into 2 different files: one including all LSVs (flag –show-all) and another file with the flag “–changing-between-group-dpsi 0.2” to select for those LSVs (Local Splice Variations) with a change in inclusion of at least 20% between pre- and post-relapse samples.

### Nanopore long-read direct RNA sequencing

From whole Raji cells, total RNA was isolated using the Maxwell® RSC simplyRNA Cells Kit (Promega). 500ng of mRNA was isolated from total RNA using Dynabeads mRNA DIRECT kit (Invitrogen) and used for direct RNA (SQK-RNA002, ONT) library preparation. Subsequently, each library was loaded into a Spot-ON Flow Cell R9 version (FLO-MIN106D, ONT) and sequenced in a MinION Mk1B device (ONT) for 48 hours. Raw Fast5 files were converted to fastq with guppy (v3.4.5), followed by alignment to the gencode version of hg38 (v30) using minimap2 (v2.18); the resulting bam file was visualized using the Integrative Genomics Viewer (v2.11.0).

### Transient expression of CD20 mRNA isoforms

The full-length sequence of MS4A1 V1 (NM_152866), V2 (NM_152867), V3 (NM_021950) and V4 were cloned into the MXS_CMV::PuroR-bGHpA plasmid, replacing the PuroR ORF in the process, to generate the V1, V2, V3 or V4 expression plasmids.

MXS_CMV::PuroR-bGHpA was a gift from Pierre Neveu (RRID:Addgene_62439) ^44^. Subsequent deletions and mutations were also generated via Gibson assembly. Plasmids were transfected into HEK293T cells using the ViaFect™ Transfection Reagent (Promega) according to manufacturer instructions ^45^.

### Stable expression of CD20 mRNA isoforms

The T2A sequence within the pCDH-EF1α-MCS-T2A-Puro plasmid (System Biosciences) was replaced with the IRES sequence from the pMXs-IRES-Puro (Cell Biolabs) to generate the pCDH-EF1α-MCS-IRES-Puro lentiviral transfer plasmid. The 5’-UTR and CDS of MS4A1 V1, V2 and V3 were cloned into the pCDH-EF1α-MCS-IRES-Puro vector to generate the V1-, V2- and V3-Puro plasmids. The CD20-targeting gRNA resistant V1-, V2-and V3-rCD20 lentiviral transfer plasmids were generated by mutating the CCTGGGGGGTCTTCTGATGATCC sequence, found within the CD20 CDS of the V1-, V2- and V3-Puro plasmids, to GTTGGGCGGACTACTTATGATTC; and replacing the puromycin resistance gene with a blasticidin resistance gene. Subsequent deletions and mutations were also generated via Gibson assembly.

### CD20 knockout

A gRNA sequence targeting the GGATCATCAGAAGACCCCCC sequence within the CD20 CDS, and a non-specific sequence targeting GTTCCGCGTTACATAACTTA, was cloned into the lentiCRISPR v2 lentiviral vector for co-expression with Cas9. The lentiCRISPR v2 plasmid was a gift from Feng Zhang (Addgene plasmid # 52961) ^46^. We stably expressed these plasmids in OCI-Ly8 cells and used fluorescence activated cell sorting (FACS) to enrich for cells that stained negative with PE anti-human CD20 antibodies (clone 2H7, Biolegend). This sorted CD20-negative cell population were the OCI-Ly8 CD20KO cells.

### Morpholino treatments

The Ex2-1, Ex2-2, and 5ex3 morpholino sequences were as follows: AGTAGAGATTTTGTTCTCTCTTGTT, ATTGTCAGTCTCTTCCCCACAGAAT, and CTGCTGAGTTCTGAGAAAGGAGATG, respectively. Standard morpholinos (Gene Tools, LLC) were electroporated into Raji and OCI-Ly8 cells using the Neon™ Transfection System 10 μL Kit (ThermoFisher). Briefly, for every 10 μL tip, half a million live cells were electroporated in buffer R containing 10 mM of the morpholino at 1350 V and 30 ms pulse width. The Random Control 25-N morpholino (Gene Tools) was electroporated as the Ctrl-PMO. Raji, OCI-Ly8 and MEC-1 cells were incubated in cell culture media containing 10 mM Vivo Morpholinos (viMO) for 3 h before the media was replaced with fresh media. In Ctrl-viMO, the reverse sequence of 5ex3, GTAGAGGAAAGAGTCTTGAGTCGTC, was used. Cells were tested for rituximab sensitivity on day 2 post morpholino treatment.

### Evaluation of rituximab sensitivity

To cell cultures containing a million live cells per mL, rituximab (Roche) was added. After 15 min, normal human serum (Complement Technology) was then added to a 20% (v/v) concentration as a source of complement, for complement-dependent-cytotoxicity. After another 2 h incubation, cell viability was evaluated by PI staining before acquisition on a BD Accuri™ C6 Cytometer (BD Biosciences), or by the WST-1 cell proliferation reagent according to the manufacturer’s protocol (Sigma-Aldrich). The WST-1 absorbance signal (A450nm – A690nm) was measured using the BioTek Synergy 2 instrument. The blank control well contains the same volume of culture medium and WST-1 but no cells. The % cell viability was calculated as (absorbance at X concentration of rituximab□−□absorbance of blank control well)/(absorbance with no rituximab – absorbance of blank control well). Dose response curves and IC50 values were calculated using the nonlinear regression curve fit function in GraphPad Prism 5 (v5.01) with the top and bottom of the curve constrained to 100 and 0% respectively.

### Manufacturing of CD20-directed human CAR T cells (CART-20)

CAR20 plasmid was custom generated by TWIST Bioscience and encompasses 1F5 scFv ^47-49^, CD8A hinge, 4-1bb costimulatory domain, and CD3ζ stimulatory domains. Healthy primary human T cells were obtained from the Human Immunology Core of the University of Pennsylvania. To produce CART20, CD4+, and CD8+ cells were combined at a 1:1 ratio and activated using anti-human CD3/CD28 Dynabeads (Invitrogen, #40203D) at a 3:1 ratio. 24 hours (Day 2) post activation, T-cells were infected with a lentiviral vector containing the CAR20 transgene (multiplicity of infection ∼1.5 viral particles / T cells). Magnetic beads were removed from T cells on day 6, and the transduction efficiency of CAR (% CAR+) was measured by flow cytometry using a PE-anti G4S antibody (Cell Signaling Technology #38907S). T-cells were expanded, counted every other day, and cryopreserved at a cell volume ≤ 350 fL. Prior to functional assays, T cells were thawed and rested overnight at 37°C.

### Mosunetuzumab cytotoxicity assays

OCI-Ly8 were transduced to express green click-beetle-luciferase and ZsGreen using pHIV-Luc-ZsGreen [a gift from Bryan Welm (Addgene plasmid # 39196)]. These cells were cultured with healthy donor T cells (E:T ratio of 5:1) and different concentrations of Mosunetuzumab (Genentech, Roche): 0, 1, 10, 100 ng/ml. After 24 hours, cancer cell survival was monitored as the luciferase bioluminesce using a BioTek Synergy H4 Imager (560 nm). Cell viability (%) was calculated relative to control cancer cells and T cells without mosunetuzumab.

### CART20 cytotoxicity assays

OCI-Ly8 were co-cultured with CART20 or untransduced T-cells at 1:4 effectors to target ratio for 24 hours. Cell survival was analyzed using bioluminescent quantification as described above. Cell viability (%) was calculated relative to control cancer cells alone.

## RESULTS

### Up to four distinct 5’-UTRs of CD20 (V1, V2, V3 and V4) can be detected and quantified via long-read direct RNA sequencing, Illumina RNA sequencing, and RT-qPCR

We and others have implicated aberrant splicing as recurrent mechanism of resistance to CD19 and CD22-directed immunotherapies [reviewed in ^50,51^]. Here, we hypothesized that the CD20-encoding gene, *MS4A1*, may also undergo alternative splicing, during which exons from the same gene are joined in different combinations, leading to the generation of different mRNA isoforms. To identify full-length cap-to-poly(A) mRNA isoforms of CD20 and rule out reverse transcription artifacts which are common in cDNA-seq approaches ^52^, we performed long-read Oxford Nanopore direct RNA sequencing on the Raji cell line. Using this approach, we found three mRNA isoforms of CD20 corresponding to annotated transcripts, dubbed variants 1-3, or V1, V2 and V3 (NCBI annotation release 109.20211119 for the *MS4A1* gene) [**Figure 1A**]. These mRNA isoforms had distinct 5’-UTRs but had identical coding sequences. We also detected a fourth, albeit rare variant, with an unannotated permutation of the 5’-UTR exons which we dubbed V4 [**Figure 1A, bottom**]. For each 5’-UTR variant, at least two mRNA isoforms existed due to two potential alternative polyadenylation sites within the 3’-UTR of CD20 [**Figure 1A**].

**Figure 1.**
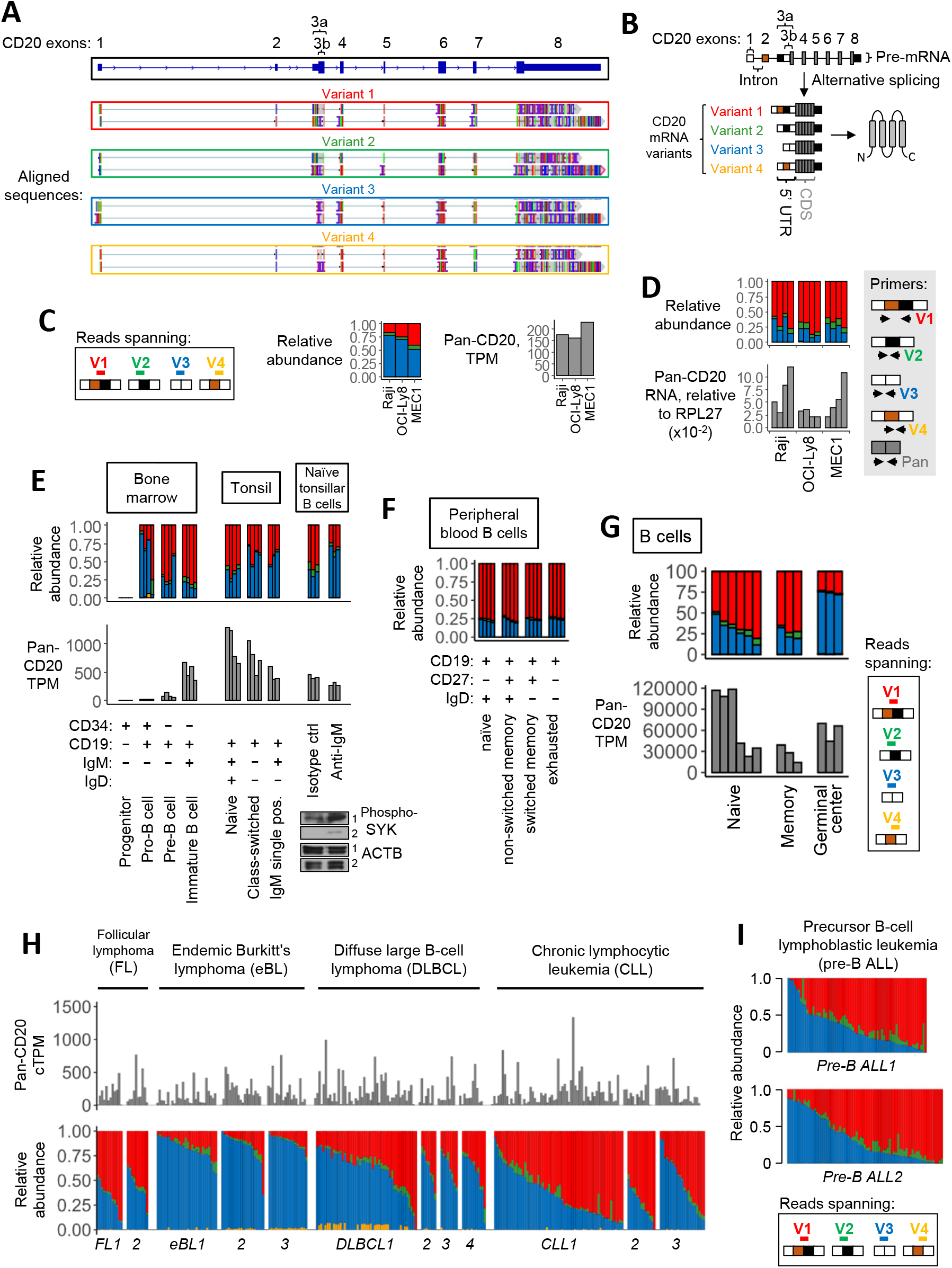
Normal and malignant B cells express four CD20 transcript variants (V1 to V4) with distinct 5’-UTRs. **(A)** Oxford Nanopore long-read direct RNA sequencing of mRNA from Raji cells. Sequence alignments corresponding to full-length, cap-to-poly(A) CD20 mRNAs with four 5’-UTR splice variants and 2 alternative 3’UTRs are shown. Alignments were extracted from and visualized in the IGV viewer. **(B)** Diagram depicting the CD20 pre-mRNA and the four distinct 5’-UTR splice variants, V1 to V4. **(C-D)** Relative abundance of V1(red), V2(green), V3(blue) and V4(yellow) isoforms in OCI-Ly8, Raji, and MEC-1 cells. Panel **(C)** is based on RNA-seq data, with the stack plot showing the ratio of sequencing reads that map to the unique exon junctions found in each 5’-UTR variant of CD20. Here and below these variants are color-coded as shown in the “Reads spanning” panel. Here and below pan-isoform reads mapping to any exon of CD20 are shown in the right for comparison as Transcripts Per Million (TPM). Panel (**D)** is based on RT-qPCR-mediated quantification of pan- and 5’-UTR variant-specific CD20 levels in OCI-Ly8, Raji, and MEC-1 cells. RNA levels are normalized to the reference gene, RPL27. Each bar represents the average from each repeated experiment (N=4). **(E-I)** Relative abundance of V1(red), V2(green), V3(blue) and V4(yellow) isoforms in primary samples corresponding to normal and malignant B cells. Each bar represents data from a single donor. The cTPM values at the top of panel (**H)** are TPM values corrected for potential batch effects between different datasets. Normal B cell subsets were FACS-enriched from the human bone marrow and tonsils **(E)** and peripheral blood **(F)**. In **(E)**, CD19+IgM+IgD+ naïve tonsillar B cells were treated with anti-IgM or an isotype control. Western blots of phosphorylated SYK are displayed at the bottom right corner as a marker of B cell activation, with actin serving as a loading control.

While 4 different 5’-UTR variants of CD20 [summarized in **Figure 1B**] were detected via long-read direct RNA sequencing, we were unable to confidently quantify the relative abundances of these variants due to the low throughput, and hence low read depth, of direct RNA sequencing. To determine the relative abundance of each 5’-UTR variant, we also analyzed available high-depth short-read RNA-seq data (>50 million reads of 2x150bp) corresponding to Raji BL, OCI-Ly8 DLBCL and MEC-1 CLL cell lines. Our custom pipeline quantified RNA-seq reads that mapped to the unique exon-exon junctions found in each of the four 5’-UTR variants (V1 to V4). In all three cell lines, V3 was the predominant isoform, making up at least 50% of all reads, followed by V1, V2 and V4 [**Figure 1C**]. To validate RNA-seq results, the different 5’-UTRs of CD20 were also quantified using isoform-specific quantitative real-time PCR (qPCR) assay as described in **supplemental Methods** and **supplemental Figure 1**. In Raji, OCI-Ly8 and MEC-1 cells, the RT-qPCR assay identified V1 and V3 as the two most abundant 5’-UTR variants [**Figure 1D**], which was consistent with our RNA-seq findings [**Figure 1C**].

### The 5’-UTRs of CD20 are alternatively spliced in normal and malignant human B cells

Having determined that our RNA-seq pipeline broadly reflects the diversity of CD20 5’-UTR isoforms, we used it to analyze several in-house ^35-37^ and publicly available ^38^ datasets. As in B-lymphoid cell lines [**Figure 1C-D**], the V1 and V3 isoforms accounted for the bulk of CD20 transcripts in both normal B cells and malignancies derived therefrom (i.e., precursor B-ALL, eBL, CLL, DLBCL, and FL). [**Figure 1E-I**]. In most normal B cell subsets from bone marrow, tonsils [**Figure 1E**] and peripheral blood [**Figure 1F**] the V1 transcripts outnumbered V3. The few exceptions included class-switched B cells from tonsils [**Figure 1E**, middle panel] and germinal center (GC) B cells [**Figure 1G**], where V3 predominated. Since BCR activation occurs in GCs prior to class-switching, we analyzed RNA-seq data from naïve tonsillar B cells that were activated with antibodies directed against IgM, as described previously ^53,54^. As expected, B cell receptor (BCR) ligation led to SYK phosphorylation and also to an increase in the relative abundance of V3 [**Figure 1E**, right panel]. These findings suggest that alternative splicing in the 5’-UTR of CD20 is tightly modulated during normal B cell development. Compared to this relatively homogenous distribution in normal B-cell samples from different donors [**Figure 1E-G**], a high degree of intertumoral variability in the relative abundance of the V1 and V3 isoforms, was observed for CLL, DLBCL, FL [**Figure 1H**] and precursor B-ALL [**Figure 1I**].

### EBV infection increases the abundance of V3 and V4

Among the B-cell neoplasms, eBLs stood out as having relatively high levels of V3, per RNA-seq [**Figure 1H, supplemental Figure 2A**]. The high V3 may be a consequence of EBV infection, as up to 95% of eBL cases are associated with this human pathogen ^55^. To test this hypothesis, we analyzed lymphoblastoid cell cultures (LCLs) derived from EBV-infected mononuclear cells. We observed a clear increase in V3 even as total CD20 mRNA levels were lower than in DLBCL and FL [**Figure 2A**]. Our RT-qPCR assays also detected a similar increase in V3 and V4 in B cells, 3 days after exposure to EBV, relative to the untreated controls [**Figure 2B**, top]. Similar results were obtained using the published RNA-seq dataset ^56^ where B cells rapidly upregulated V3 and V4 as early as one day after exposure to EBV [**supplemental Figure 2B**]. Thus, alternative splicing in the 5’-UTR of CD20 is affected by EBV infection.

**Figure 2.**
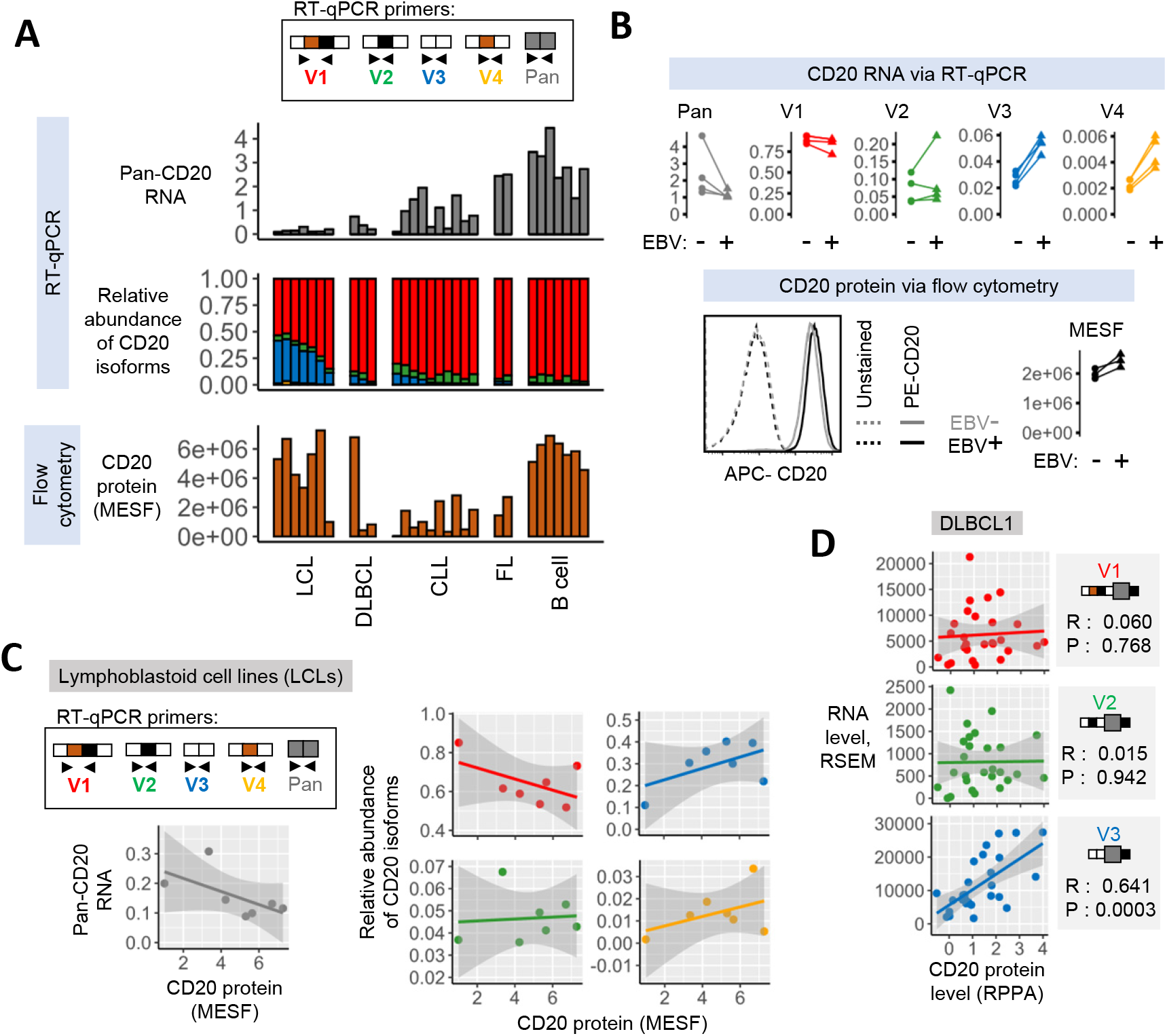
CD20 protein levels positively correlate with the abundance of V3 and V4 but not V1, V2 or pan-isoform CD20 mRNA. **(A-C)** The levels of pan-isoform and 5’-UTR variant-specific mRNAs and CD20 protein (expressed as Molecules of Soluble Fluorochrome or MESF) in primary or immortalized human samples. They were quantified using RT-qPCR and flow cytometry, respectively. RNA levels are normalized to three reference genes: RPL27, ACTB and GAPDH. Each bar or point represents data from a single donor. Panel **(A)** displays data from EBV-transformed lymphoblastoid B-cell lines (LCLs), peripheral blood B cells of healthy donors, and patients with DLBCL, CLL or FL. In **(B)**, peripheral blood B cells from four different donors (N=4) were cultured with the B95.8 strain of EBV (EBV+) or without EBV (EBV-) for 3 days prior to CD20 expression quantification. **(C)** Regression analysis of 7 LCL samples from panel **(A)** where CD20 protein levels are independently correlated to each of the known CD20 mRNA isoforms. **(D)** Similar regression analysis of 27 DLBCL samples from the TCGA consortium for which Reverse Phase Protein Array (RPPA) and RNA-seq data are available. The shaded area around the regression line indicates the standard error. Spearman’s rank correlation coefficient, R, and the two-tailed p-value, P, as calculated in MATLAB is shown.

### An increase in V3 and V4 coincides with elevated CD20 protein

The increase in V3 and V4 after EBV infection of B cells [**supplemental Figure 2B, Figure 2B**, top] was notable as it occurred in tandem with elevated levels of cell-surface CD20, as measured by flow cytometry. EBV infection increased CD20 protein levels despite also down-regulating total CD20 mRNA, as observed in our experiments [**Figure 2B**] and also as described in the literature ^56^. This trend was preserved in LCLs, where elevated levels of V3 coincided with elevated levels of surface CD20 protein, relative to CLL, DLBCL and FL. In contrast, pan-isoform CD20 mRNA levels did not correlate with CD20 protein levels as its levels in LCLs were among the lowest [**Figure 2A**]. When we compared individual LCL samples, we again found that CD20 protein levels did not positively correlate with the levels of the V1, V2, or total CD20 mRNA, and instead correlated with the levels of V3 or V4 mRNA isoforms [**Figure 2C**]. The same correlations were found in the DLBCL samples from the TCGA consortium, which had Reverse Phase Protein Array (RPPA)-based measures of CD20 protein [**Figure 2D**]. Taken together, these findings suggest that CD20 protein levels are controlled exclusively by the abundant V3 isoform.

### The extended 5’-UTRs variants (V1 and V2) have increased transcript half-life but reduced ribosome recruitment

Our CD20 data mirrored a similar expression pattern established by us for CD22, whereby its protein levels positively correlated with the abundance of productive CD22 mRNA isoforms but not with total CD22 mRNA (which included a large fraction of AUG-lacking non-coding transcripts ^37^. This led us to hypothesize that a similar mechanism was at play for CD20. While the V1 and V3 CD20 mRNA isoforms have identical coding sequences they possess distinct 5’-UTRs which can affect CD20 protein production by altering mRNA turnover and/or translation rates. To investigate if there was a difference in mRNA turnover, we used our isoform-specific RT-qPCR assay to measure the rate of turnover or decay for each CD20 mRNA isoform in OCI-Ly8 and Raji cells after inhibiting transcription in these cells with Actinomycin D. Relative to V1 and V2, mRNA decay rate for V3 and V4 very slightly higher [**Supplemental Figure 3**]. The RT-qPCR assay also allowed us to measure the rate of translation for each 5’-UTR isoform in Raji and OCI-Ly8 cells using polysome profiling as described previously ^57,58^. Relative to V3 and V4, a smaller fraction of V1 and V2 molecules was detected in the high-density sucrose gradient fractions [**Figure 3A**]. This pattern is indicative of a lower rate of translation for V1 and V2 relative to V3 and V4.

**Figure 3.**
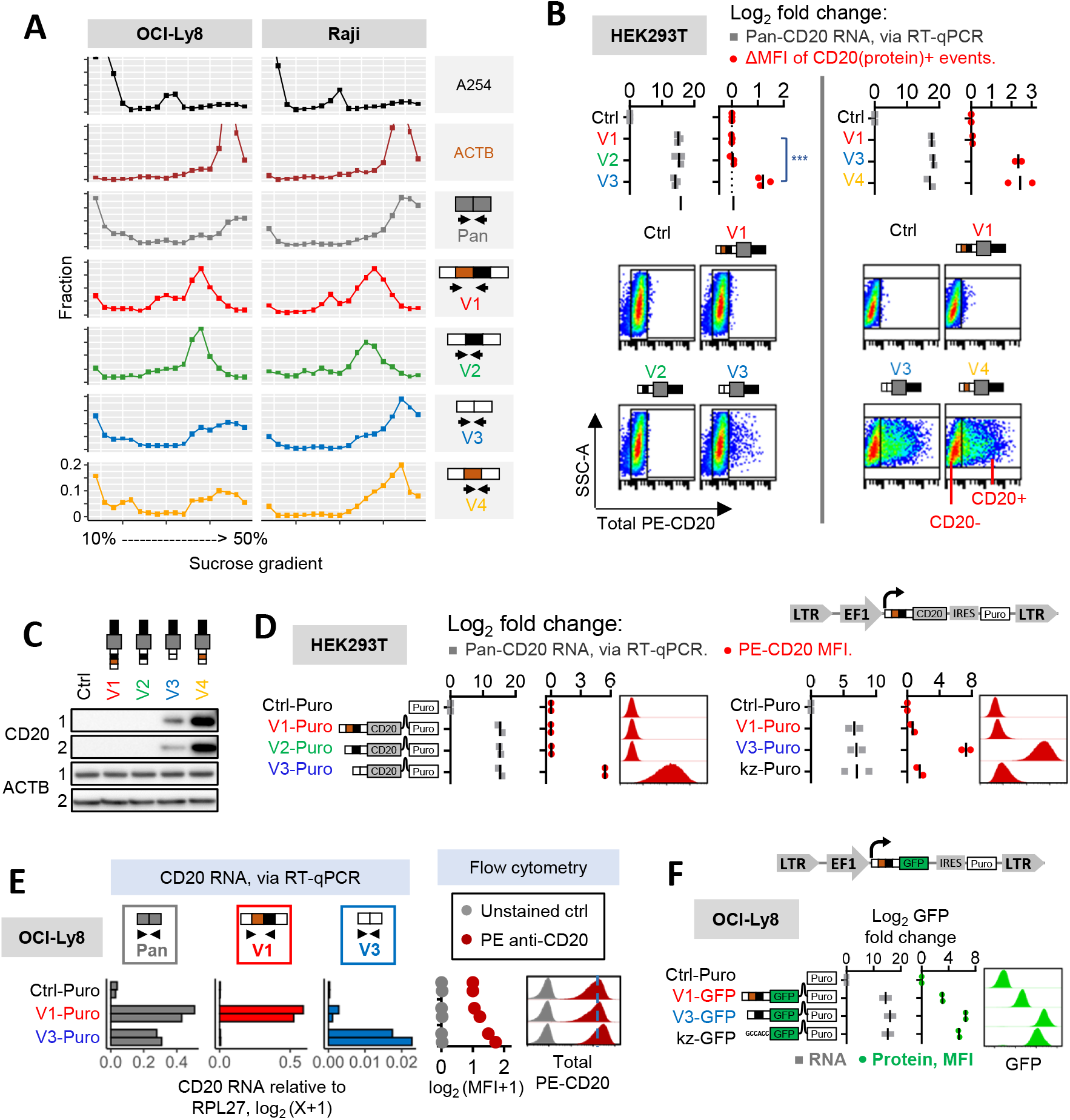
CD20 mRNA isoforms V3 and V4, but not V1 and V2, are efficiently translated into protein. **(A)** Polysome profiling of OCI-Ly8 and Raji cells. The top panel shows the ribosomal content measured at absorbance 254□nm, while the bottom panels show the relative distribution of specific transcripts across the sucrose gradient fractions, as measured via RT-qPCR [see Supplemental Methods]. **(B)** CD20 mRNA and protein measurements in HEK293T cells transiently transfected to express V1, V2, V3 or V4 mRNA isoforms. Pan-isoform CD20 transcripts were measured using RT-qPCR. Total CD20 protein was measured with flow cytometry and expressed as the delta median fluorescence intensity (ΔMFI) [see Supplemental Methods]. Representative scatter plots are shown in the bottom half. The *** symbols indicate P<0.001 per 1 way ANOVA test (pwc: Bonferonni). **(C)** Immunoblotting analysis of CD20 protein levels in cells from the previous panel. Β -actin (ACTB) served as the loading control. Data from 2 independent experiments are shown. **(D)** CD20 mRNA and protein measurements in HEK293T stably expressing the empty vector (Ctrl-Puro) or the 5’-UTR and coding sequences of V1, V2 or V3 (V1-, V2-or V3-Puro). The kz-Puro control has the “GCCACC” Kozak consensus as its sole 5’-UTR element. The levels of CD20 mRNA were quantified with RT-qPCR. The median fluorescence intensities (MFI) of total cellular PE-CD20 were determined by flow cytometry. Representative histograms are shown. **(E)** The same experiment performed on OCI-Ly8 cells. **(F)** CD20 mRNA and protein measurements in OCI-Ly8 cells stably expressing the V1 or V3 5’-UTR sequences followed by a GFP reporter (V1- and V3-GFP). The kz-GFP control has the “GCCACC” Kozak consensus as its sole 5’-UTR element. GFP mRNA levels were measured using RT-qPCR. The median fluorescence intensity (MFI) of GFP was measured using flow cytometry. Representative histograms are shown. In **(B, D-F)**, RNA levels are normalized to the reference gene, RPL27. Each bar or dot represents the average from repeated experiments (N≥2). RNA and MFI values are expressed as log2 fold change relative to the Ctrl or Ctrl-Puro sample, as indicated.

### Heterologous expression of V3, but not V1, enhances CD20 protein expression

Inefficient translation of V1 and V2 transcripts could be rate-limiting for CD20 protein levels. To investigate this scenario, we engineered HEK293T and OCI-Ly8 cells to express each 5’-UTR variant (V1 through V4) of CD20, and then investigated whether these cells had increased levels of CD20 mRNA (measured via RT-qPCR) and protein (measured via flow cytometry and/or western blotting), relative to control cells transfected with empty vector (Ctrl or Ctrl-Puro). In HEK293T cells, which do not express CD20 endogenously, transient transfection with full-length V1, V2, V3 and V4 cassettes led to uniformly elevated levels of CD20 mRNA [**Figure 3B**, top].

However, these increases in mRNA levels were accompanied by increased CD20 protein production only in V3- and V4-transfected cells, while V1- and V2-transfected cells remained CD20 protein-negative and were indistinguishable from the Ctrl cells [**Figure 3B-C**]. This 5’-UTR-specific difference in CD20 protein levels was independent of the 3’ UTR since cells transfected with V3:del-3’ (the 3’ UTR-deletion mutant version of V3) still had markedly more CD20 protein staining than those transfected with V1:del-3’ or V2:del-3’ [**supplemental Figure 4A**]. In these and subsequent experiments, the GCCACC Kozak consensus (“kz”) alone was used as a positive control 5’-UTR element **[supplemental Figure 4B]**.

Similar results were obtained when HEK293T or OCI-Ly8 cells were stably transduced with lentiviral vectors to express these variants lacking the 3’ UTRs (V1-Puro, V3-Puro, etc.) [**Figure 3D,E**]. Unlike HEK293T cells, OCI-Ly8 cells express endogenous CD20 protein which we detected in the empty vector-transduced cells(Ctrl-Puro), and only transduction with V3-Puro, but not V1-Puro, increased total CD20 protein levels [**Figure 3E**, right]. To determine whether 5’-UTRs alone are responsible for the difference in expression, we generated GFP reporters lacking any CD20 coding sequences. Once again, only V3 and not V1 constructs yielded robust expression of the reporter gene, as evidenced by flow cytometry [**Figure 3F**].

### The extended 5’-UTRs (V1 and V2) contain uORFs and an RNA stem-loop structure that repress translation

The inefficient translation of V1 and V2 implied the presence of repressive 5’-UTR elements. We reasoned that such elements would localize in exon 3a based on two observations. Firstly, exon 3a was included in V1 and V2 but not in V3 and V4. Secondly, a truncated mutant construct which retained exon 3a but lacked exons 1 and 2 (i.e., Δex1-2) still retained the ability to repress CD20 protein levels [**supplemental Figure 4C**]. Within the exon 3a sequence, we found multiple open reading frames (uORFs). At the exon 3a-b boundary immediately upstream of the CD20 start codon, a stem-loop secondary structure is predicted by the RNAfold algorithm ^59^ [**Figure 4A**]. Since translational repression by uORFs and stem-loops has been observed in other genes ^60,61^, we hypothesized that CD20 could be regulated in the same manner. To investigate this possibility, we mutated all the start codons of the uORFs in V1 from AUGs to AUCs to generate V1AUC-Puro. We also generated the V1DelStem-Puro variant with disrupted base pairing in the stem-loop, alone or in combination with the AUC variant, and the V1Stem-Puro, where the hairpin was stabilized via additional base pairing [**Figure 4B]**. We then measured CD20 RNA and protein levels by RT-qPCR and flow cytometry, respectively, compared to the Ctrl-Puro empty vector

We found that V1AUC-Puro cells remained CD20 protein-negative [**Figure 4C**], as were V1Stem-Puro cells [**Figure 4D**]. There was a minor recovery in CD20 protein levels with the V1DelStem-Puro variant [**Figure 4E**]. Notably, disrupting both the uORFs and the stem-loop (V1ATCDelStem-Puro) led to a near complete recovery in CD20 protein levels [**Figure 4F]**. These results suggest that these two elements are responsible for the negligible CD20 protein being produced from V1 and V2.

**Figure 4.**
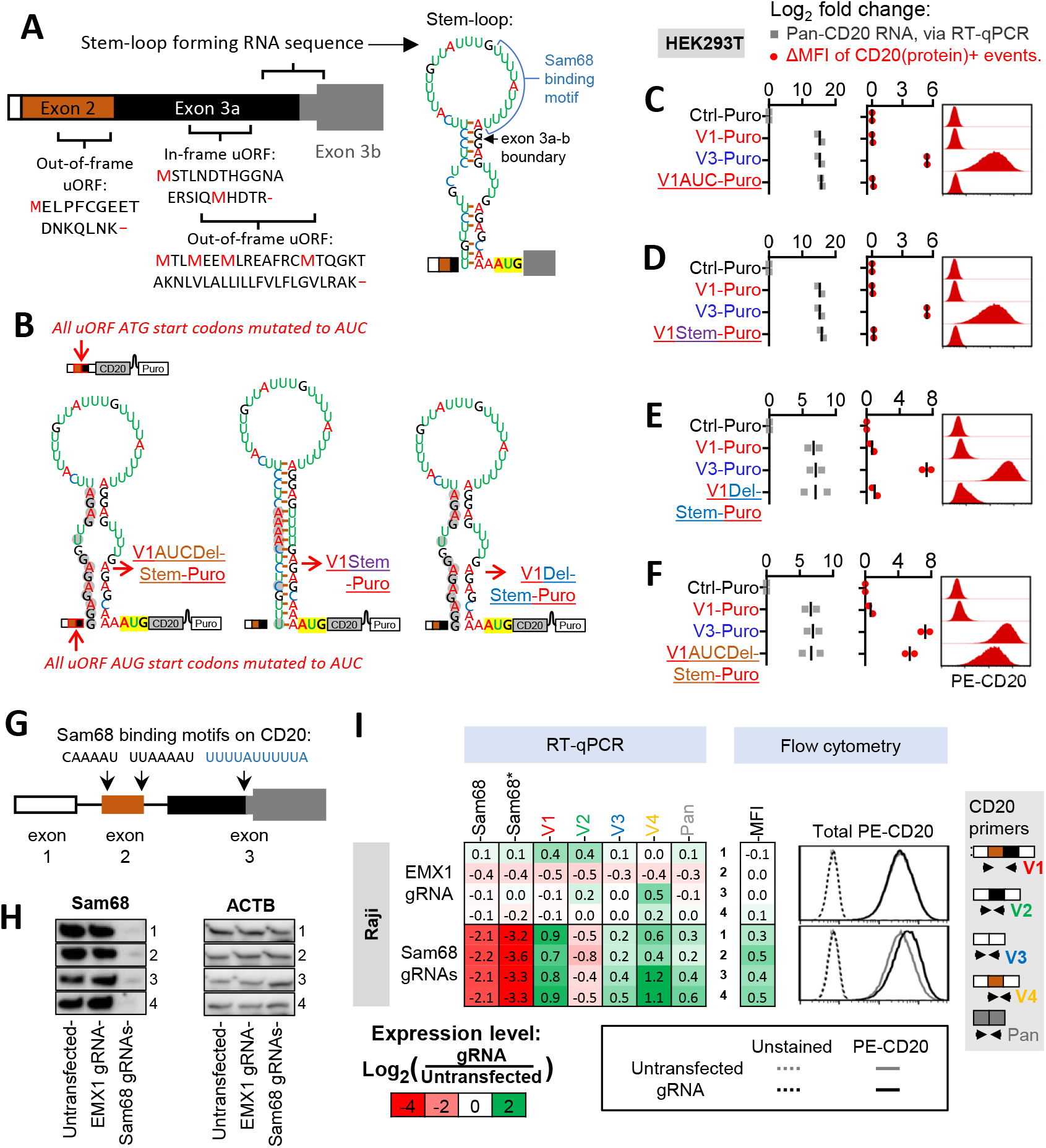
An RNA stem-loop structure and uORFs repress translation of the CD20 V1 isoform. **(A)** Diagram depicting the uORFs and the stem-loop predicted by the RNAfold web server to form in the 5’-UTR of the CD20 V1 isoform. **(B)** Diagrams of mRNA product corresponding to various derivatives of V1-Puro. All AUG start codons found in the 5’-UTR of V1-Puro were mutated to AUC in the V1AUC-Puro and V1AUCDel-Stem-Puro constructs. “Stem” and “Del-Stem” refer to mutations stabilizing and destabilizing the predicted secondary structure, respectively. **(C-F)** Expression of V1- and V3-Puro and the V1-Puro mutants in stably transduced HEK293T cells. Pan-isoform CD20 RNA levels were quantified using RT-qPCR. The median fluorescence intensity (MFI) of total cellular CD20 protein was measured using flow cytometry. Representative histograms are shown on the right. Each bar or dot represents the average from repeated experiments (N=2). All values are expressed as log2 fold change relative to the Ctrl-Puro sample. **(G)** Diagram depicting the putative Sam68 binding sites found within exons 2 and 3 of V1. **(H-I)** Sam68 and CD20 expression measurements in Raji cells electroporated with a mixture of two Cas9-gRNA ribonucleoproteins targeting the CDS of Sam68. Shown in **(H)** is immunoblotting analysis of Sam68 protein expression, with ACTB serving as the loading control. Shown on the left side of **(I)** are Sam68 and CD20 mRNA levels, as quantified using RT-qPCR. Two PCR primers pairs measured non-overlapping regions within the Sam68 transcript. The second primer pair is denoted with an asterisk (*). Shown on the right side of **(I)** are median fluorescence intensities (MFI) of total CD20 protein, as determined using flow cytometry. Representative histograms are shown on the far right. In both **(H)** and **(I)**, each row in the immunoblotting image and the heatmap represent an independent experiment (N=4).

### Sam68 contributes to suppressed CD20 protein levels

We searched for RNA-binding proteins (RBPs) that could be responsible for often high V1-to-V3 ratios and translational repression of CD20. Using the SpliceAid algorithm ^62^, we identified within exon 2 [**Figure 4G**] and notably within the loop structure of exon 3a [**Figure 4A**] several putative binding sites for Sam68 (aka KHDRBS1) a splicing factor known to bind to AU-rich sequences ^63^. To determine if Sam68 regulates CD20 splicing, we electroporated Raji cells with a mixture of two Cas9-gRNA ribonucleoproteins targeting the coding sequence (CDS) of Sam68. Relative to control gRNAs targeting the irrelevant EMX1 locus and untransfected control cells, the Sam68 gRNAs successfully reduced Sam68 protein levels [Western blots in **Figure 4H**]. This was predictably accompanied by a reduction in Sam68 mRNA levels as measured by RT-qPCR [heatmap in **Figure 4I**, two leftmost columns]. We also observed a re-distribution of all 4 5’-UTR variants: a reduction in V2 mRNA levels and increases in V1, V3, and V4, and total CD20 mRNA[**Figure 4I**, five rightmost columns]. We also observed a modest increase in CD20 protein levels [flow cytometry in **Figure 4I**, right]. The same shifts in CD20 mRNA splicing and protein levels were evident when we stably knocked down Sam68 in Raji and OCI-Ly8 cells with two different shRNAs [**supplemental Figure 5A, B**]. Taken together, our findings suggest that Sam68 is at least partly responsible for the selection of CD20 5’-UTR isoforms and for suppressed CD20 protein levels. However, we could not distinguish between its effects on splicing and on overall mRNA levels.

**Figure 5.**
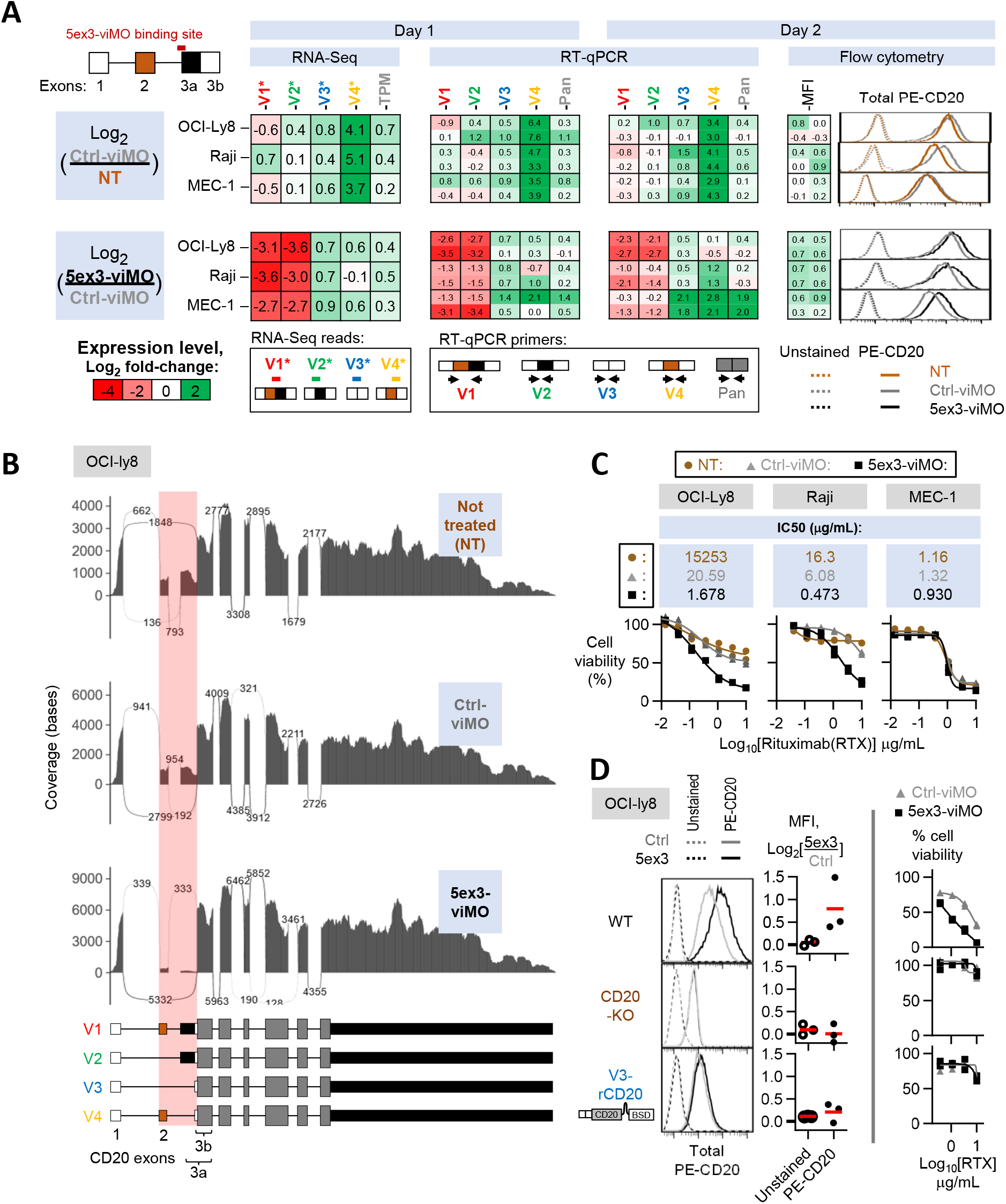
Shifting CD20 splicing towards V3 and V4 increases CD20 protein expression and rituximab-mediated cytotoxicity. **(A-C)** CD20 mRNA and protein measurements in OCI-Ly8, Raji and MEC-1 cells treated with either the solvent control (NT), Ctrl-viMO, or 5ex3-viMO. **(D)** The same experiment performed on OCI-Ly8 variants with the intact *MS4A1* gene (“WT”, top), *MS4A1* gene knocked out (“CD20-KO”, middle), or knocked-out *MS4A1* gene replaced with the V3 expression cassette (“V3-rCD20”, bottom). CD20 RNA levels were quantified using RNA-seq [leftmost heatmap in **(A)** and sashimi plot in **(B)**] and RT-qPCR [middle heatmaps in **(A)**]. For the RNA-seq data in **(A)**, V1*-4* are reads that mapped to the unique exon junctions found in each 5’-UTR variant of CD20, color-coded as shown in the “RNA-seq reads” panel. The median fluorescence intensity (MFI) of total CD20 protein were determined using cytometry [rightmost heat map in **(A)**, top panel in **(C)**, and the left and middle panels in **(D)**]. Representative histograms are shown on the far right of **(A)** and far left of **(D)**. In the bottom of **(C)** and the right panel of **(D)** cells were additionally treated with increasing concentrations of rituximab (RTX), and cell viability (plotted) was measured using the WST-1 assay. All values are normalized to the “no rituximab control”. In **(A, C, D)**, each dot in a graph or value in a heatmap represents independent experiments (N≥2), with mean value indicated by red lines in the graphs.

### Morpholinos redirect splicing towards V3 and V4 to augment rituximab-mediated cytotoxicity

To modulate CD20 splicing more directly, we tested three different phosphorodiamidate Morpholino oligomers (PMO) ^64^ sequences that target exons 2-3 in CD20 pre-mRNA [**supplemental Figure 5C**, top right]. These PMOs were electroporated into Raji and OCI-Ly8 cells. Among the PMOs tested against a non-specific control sequence pool (Ctrl-PMO), the “5ex3-PMO” sequence (targeting the junction between CD20 intron 2 and exon 3a) was the most effective at reducing V1 and V2 mRNA while increasing V3 and in particular V4 mRNA (∼3-fold change, as measured by RT-qPCR). It was also superior to Ex2-1 and Ex2-2 at increasing CD20 protein levels, as measured by flow cytometry) [**supplemental Figure 5C**].

Next, we obtained 5ex3 in the Vivo-Morpholino (viMO) format [**Figure 5A**, top left], where it could be added directly to culture media for efficient delivery into cells ^65^. We tested 5ex3-viMO on OCI-Ly8, Raji and MEC-1 cell cultures and measured the levels of CD20 mRNA (by RT-qPCR and RNA-seq) and total cellular CD20 protein (by flow cytometry). As a control, we used the oligomer with the reverse order of 5ex3 nucleotides (Ctrl-viMO), which did little to affect V1, V2, and V3 variants, but induced non-specific increases in V4 and CD20 protein levels relative to the no-Morpholino treatment (NT) cells [**Figure 5A**, top rows]. To account for these sequence-independent effects, we normalized the 5ex3-viMO treated samples to the Ctrl-viMO treatment. Even after normalization, 5ex3-viMO treatment induced clear shifts in CD20 splicing, namely reducing V1 and V2 mRNA levels while increasing V3 and V4 variants [**Figure 5A**, bottom rows]. Visual analysis of the sashimi plots corresponding to the RNA-seq data confirmed that the 5ex3-viMO had a noticeably greater effect on reducing exon 2 and 3a usage [**Figure 5B**, exons within the red boxes], without affecting the usage of the other exons [**Figure 5B**, exons outside of the red boxes]. Most importantly, relative to the Ctrl-viMO treatment, 5ex3-viMO-treated cells had elevated CD20 protein staining [**Figure 5A**, bottom right corner] across multiple experiments [**supplemental Figure 5D**].

To determine whether this increase is therapeutically relevant, we adopted an anti-CD20 antibody-mediated cytotoxicity assay. In a pilot experiment, we treated OCI-Ly8 cells overexpressing Ctrl-, V1-, and V3-Puro with a wide range of rituximab concentrations and measured cell viability. We achieved 50% killing of cells expressing the V3 isoform with 4.3 µg/mL of rituximab, but failed to reach IC50 for their Ctrl and V1 counterparts [**supplemental Figure 6A**]. We then repeated the assay using OCI-Ly8, Raji, and MEC-1cells treated with Ctrl or 5ex3-viMO. In 2 out of 3 lines (OCI-Ly8 and Raji), we observed a >10-fold difference in IC50 between Ctrl- and 5ex3-viMO, as measured using the WST-1 cell viability reagent [**Figure 5C**]. This difference in sensitivity was further verified by propidium iodide (PI) assays, where cell death rather than cell survival was measured [**supplemental Figure 6B**].

To confirm that the 5ex3-viMO effect is dependent on CD20 splicing, we generated *MS4A1* knockout OCI-Ly8 cells (CD20KO) that unlike parental cells [**Figure 5D**, top row] lacked CD20 expression and were insensitive to rituximab [**Figure 5D**, middle row]. We then reconstituted them with the intron-less V3-rCD20 cassette. This re-sensitized the cells to rituximab but also made them refractory to 5ex3-viMO [**Figure 5D**, bottom row], attesting to the specificity of this reagent and to its potential clinical utility.

### V1 generates sufficient CD20 protein levels to trigger killing by CAR T cells but not mosunetuzumab

Our RNA-seq analysis of FL, DLBCL, CLL and Pre-B ALL identified a subset of samples where the entire CD20 mRNA pool consisted almost entirely of V1, with little to no V3 [**Figure 1H,I**]. To model B-cell neoplasms with such lopsided CD20 splicing pattern and to test their responses to CD20-directed immunotherapeutics, we generated an OCI-Ly8 derivative that simultaneously expressed the ZsGreen and Luciferase reporters. These cells were then transduced with *MS4A1*-specific CRISPR-Cas9 lentiviral vectors (lentiCRISPR v2) to knock out endogenous CD20 expression before being reconstituted with the V1-, V2-or V3-rCD20 constructs or an empty vector control (CD20KO). A separate batch of parental cells were transduced with the lentiCRISPR v2 vector containing a non-specific gRNA; these “WT” cells retained the expression of endogenous CD20 [**Figure 6A**]. We then evaluated the efficiency of CD20 knockouts and reconstitutions via isoform-specific CD20 RT-qPCR [**Figure 6B**] and flow cytometry for cell-surface and total CD20 protein [**Figure 6C**]. Interestingly, in the RT-qPCR assay, CD20KO cells showed roughly the same CD20 transcript levels as the WT control cells [**Figure 6B**], but these transcripts were apparently non-coding as the CD20KO cells were completely CD20 protein-negative [**Figure 6C**].

**Figure 6.**
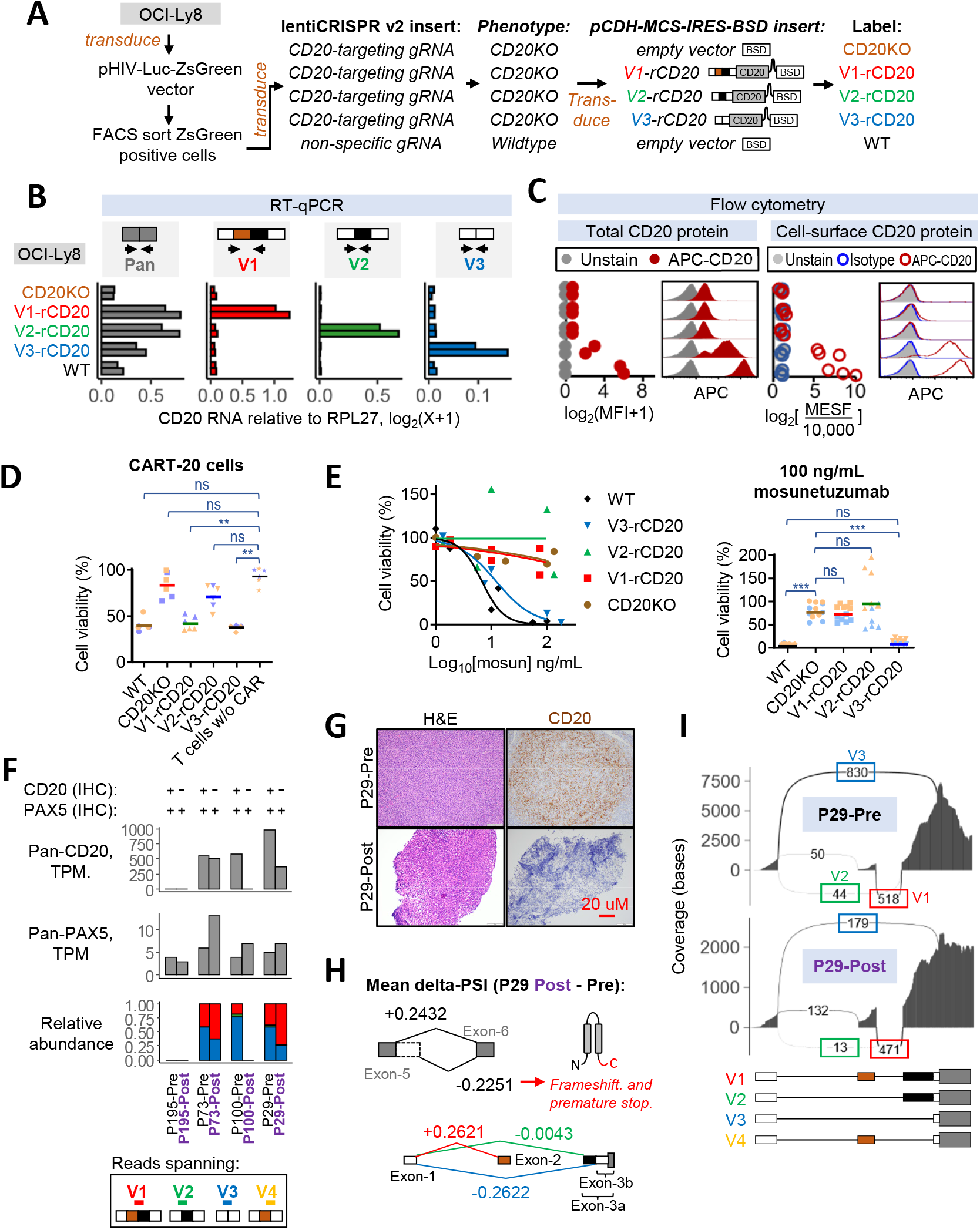
Shifting CD20 splicing from V3-to-V1 increases resistance to mosunetuzumab but not CART-20. **(A)** Diagram depicting the lentiviral vector transductions performed on OCI-Ly8 cells expressing a luciferase reporter. The *MS4A1* gene was knocked out (CD20KO) and reconstituted with V1-, V2-or V3-rCD20. The “resistant” CD20 (or rCD20) constructs have silent mutations in their coding sequence to avoid recognition by the CD20-targeting gRNA used in the knockout. **(B)** Isoform-specific RT-qPCR measurements of CD20 mRNA isoform relative to the RPL27 reference gene. **(C)** Measurement of CD20 protein levels by flow cytometry. On the left, it is quantitated as log2 fold change in median fluorescence intensity (MFI) of total APC-CD20 relative to the unstained controls. On the right, it is quantitated as cell surface CD20 levels expressed as Molecules of Soluble Fluorochrome (MESF). Each spot is an independent experiment. Representative histograms are included in each sub-panel. **(D)** Viability of OCI-Ly8 cells after 24 hours of co-culturing with CART-20 or untransduced donor T cells at 1:4 effector-to-target ratio. Cell viability (in %) is shown relative to control OCI-Ly8 cells that were not co-cultured with either CART-20 or donor T cells. **(E)** OCI-Ly8 cell viability after 24 hours of co-culturing with donor T cells and indicated concentrations of mosunetuzumab. Cell viability (in %) is shown relative to the “no mosunetuzumab” control. For **(D)**, data from two independent experiments using T cells from a single donor are shown. For **(E)**, data from four independent experiments from two unrelated donors are shown. In **(D)** and in the right panel of **(E)**, each spot represents the results from replicate wells, with different colors denoting independent experiments. The horizontal bars represent the means for all replicates. The ** and *** symbols indicate P<0.01 and P<0.001 per Kruskal-Wallis test (pwc: Dunn test). **(F)** CD20 and PAX5 status per IHC [Top] and transcript read abundance per RNA-seq [bar graphs] in paired pre- and post-mosunetuzumab follicular lymphoma (FL). Transcripts-per-million (TPM) values quantify RNA-seq reads mapping to any exon in CD20 and PAX5. The relative abundance of V1(red), V2(green), V3(blue) and V4(yellow) is the ratio of sequencing reads mapping to the unique exon-exon junctions found in each 5’-UTR variant of CD20, as color-coded in the “Reads spanning” panel. **(G)** Micrographs corresponding to P29 Pre- and Post-mosunetuzumab FL samples. FFPE sections were IHC stained for CD20 (brown colorimetric detection with DAB) with hematoxylin nuclear counterstain (blue). **(H)** Changes in percent spliced-in values (delta-PSI) or the change in ratio between reads including or excluding CD20 exons in the P29-Post sample relative to the -Pre sample. The RNA-seq data was analyzed using the MAJIQ algorithm for all possible splicing changes in CD20, but only delta-PSI values above or below 0.05 are shown. **(I)** Sashimi plots depicting density of exon-including and exon-skipping reads in the RNA-seq data corresponding to P29-Pre and -Post samples.

Furthermore, relative to WT cells, the V1-, V2- and V3-rCD20 cells had elevated levels of the cognate 5’ UTR isoforms [**Figure 6B**]. These cells could thus act as models for the complete shift in CD20 splicing towards a specific 5’-UTR variant. As expected, only reconstitution with V3-rCD20 led to a recovery in CD20 protein expression, while it remained undetectable by flow cytometry in V1- and V2-rCD20 cells [**Figure 6C**].

We then performed *in vitro* assays using 1F5 single chain variable fragment-based CAR T cells (CART-20), as described in our earlier studies on B-ALL ^37,66^. As expected, CART-20 spared CD20KO and V2-rCD20-expressing cells. Surprisingly, CART-20 was able to kill V3-rCD20- and V1-rCD20-expressing cells equally well, despite the apparent lack of detectable CD20 expression in the latter [**Figure 6D**]. We also tested the sensitivity of these cells to the bispecific CD3/CD20 antibody mosunetuzumab in the presence of donor T cells. We found that unlike CART-20, mosunetuzumab was only effective against V3-rCD20- and spared V1-rCD20-expressing cells just as it did CD20KO cells [**Figure 6E, left**]. This difference was statistically significant across two independent experiments with multiple technical replicates [**Figure 6E, right**]. Taken together, our in vitro results suggested that B cell neoplasms where the V1 variant predominates could still generate sufficient levels of CD20 protein to trigger CAR T cell-mediated cytotoxicity, but these levels would be insufficient for mosunetuzumab to be effective. This implied that the V3-to-V1 shift could underlie resistance to mosunetuzumab in patients.

### CD20 antigen loss in follicular lymphoma following mosunetuzumab treatment coincides with a V3-to-V1 shift

It was previously found that in 3/4 of the mosunetuzumab-resistant B-NHL tumors that were CD20 protein-negative by immunohistochemistry (IHC), whole exome (WES) and RNA-seq failed to detect any *MS4A1* genetic variants or loss of CD20 mRNA, which could have explained the apparent loss of CD20 protein ^16^. For this study, we obtained formalin-fixed paraffin-embedded (FFPE) tumor samples from a similar cohort of four additional patients relapsing after mosunetuzumab, and performed RNA-seq to look for changes in the splicing pattern of CD20. All four patients had paired pretreatment (“pre”) tumor samples that were CD20 positive by IHC and mosunetuzumab relapse (“post”) samples that had lost CD20 protein expression. In 3 out of 4 of the post samples, the loss in CD20 protein expression coincided with either the complete loss of CD20 mRNA (for P195 and P100) [**Figure 6F**] or an out-of-frame mutation (for P73) [**supplemental Figure 7**]. The P29-Post sample, on the other hand, retained 300 transcripts per million (TPM) of CD20 mRNA [**Figure 6F**] which contained no deleterious mutations that could explain the complete lack of CD20 protein immunoreactivity [**Figure 6G**]. Using the previously developed MAJIQ package ^67^, the RNA-seq data of P29 was analyzed for any splicing changes that could explain the loss of CD20 protein. Changes in splicing of the P29-Post sample relative to the -Pre sample was quantified as delta percent-spliced-in (ΔPSI) values. Predictably, no significant changes were found to affect the control CD19 mRNA, as no loss of CD19 expression was observed, consistent with data in the literature ^68^. Within the CD20 transcript, only two alternative splicing events exceeded a ΔPSI value of +/-0.05. The first event corresponded to a reduction (ΔPSI = -0.2251) in the usage of an alternate donor site which extended exon 5 [**Figure 6H**, top]. Since the alternate donor site leads to a frameshift and premature stop codon, the reduction in its usage is expected to increase the expression of full-length CD20 protein and would not explain the loss of CD20 expression. The second event corresponded to a decrease in splicing from exon 1 to 3b (ΔPSI = -0.2622) that occurs during the V3-to-V1 shift [diagram in **Figure 6H, bottom, and sashimi plots in Figure 6I**]. A similar shift was observed in the P73 relapse [**Figure 6F**], where it could have predated the inactivating mutation. Thus, the loss of CD20 protein in the post-mosunetuzumab P29 sample coincided with a shift in CD20 splicing away from productive V3 to translation-deficient V1.

## DISCUSSION

Here, we demonstrate that in normal and malignant B-cells, human CD20 mRNA undergoes alternative splicing to generate up to four distinct 5’-UTRs variants, V1 through V4 [**Figure 1B**]. Among them, V1 and V3 are the most abundant and together comprise more than 90% of the total CD20 transcript pool in most cell lines and primary samples [**Figure 1**]. Even though V1 and V3 contain the same open reading frame, only V3 is efficiently translated, accounting for the bulk of CD20 protein. In contrast, V1 generates negligible amounts of CD20 protein due to translational inhibition by a prominent stem-loop and multiple uORFs [**Figure 2**,**3**].

We also found that by shifting splicing away from V1 and towards V3, B cells undergoing GC reactions and class-switching could sustain or even enhance CD20 protein levels despite the reductions in total CD20 mRNA [**Figure 1F**,**G**]. This model agrees with multiple reports of elevated CD20 protein in GC cells relative to memory and naïve B cells ^69,70^. We further show that this V1-to-V3 shift is mediated at least partly by B-cell receptor signaling, since activating this pathway in tonsillar B cells by ligating surface IgM is sufficient to shift splicing toward V3 [**Figure 1E**]. Increases in the V3 isoform and CD20 protein can also be observed during EBV infection [**Figure 2B**], perhaps as part of the germinal center-like differentiation program that occurs in EBV-infected naïve B cells ^56^.

Of relevance to cancer, V3 upregulation persists after peripheral blood B cells have fully transformed into LCLs [**Figure 2A**]. The same appears to be the case in endemic Burkitt’s lymphomas (eBL) [**Figure 1H**], where 95% of cases are associated with EBV infection ^55^. Outside eBL, both V1 and V3 can be detected in most precursor B-ALL and mature B-cell neoplasms (e.g., DLBCL) [**Figure 1H**], but CD20 protein levels correlate solely with the V3 isoform [**Figure 2**]. Conversely, a subset of neoplasms expresses predominantly V1 at diagnosis, which could account for inferior responses of CLL^11-14,27^ and precursor B-ALL ^71^ to existing anti-CD20 antibody-based therapies. This suggests that the bulk of CD20 protein is produced by V3, and that in cells with active CD20 transcription the V3-to-V1 ratio is a key determinant of CD20 positivity. It remains to be determined whether the re-direction of splicing towards V3 mediates enhanced CD20 expression and rituximab sensitivity observed *in vitro* following treatment with IL4, protein kinase C (PKC) agonists, histone deacetylase inhibitors (HDACi), or the knockdown of the FOXO1 transcription factor ^72-75^.

To determine whether the V3-to-V1 shift is a relevant mechanism of resistance to CD20-directed immunotherapies, we generated OCI-Ly8 variants that express either V1 or V3 individually, and tested their responses to CART-20 and mosunetuzumab. We found that V1-expressing cells had no CD20 protein detectable by flow cytometry [**Figure 6C**], and it took the much more sensitive 5’-UTR-GFP reporter assay to observe baseline translation from the V1 mRNA [**Figure 3F**]. Nevertheless, V1-expressing cells remain sensitive to CART-20 [**Figure 6D**]. While unexpected, this finding is in agreement with the large body of literature attesting to the unique efficacy of CAR T cells against lymphoid malignancies with very low expression of target antigens ^76^. This situation might contrast with the use of rituximab and bispecific CD3/CD20 T cell engagers such as mosunetuzumab, which in our experimental system required high levels of CD20 expression only afforded by the V3 cassette [**Figure 6E**]. It then comes as no surprise that the V3-to-V1 shift was apparent in two post-mosunetuzumab follicular lymphoma relapses, P73 and P29 [**Figure 6F**]. Moreover, in the latter case, the shift was the only molecular event known to reduce CD20 expression, as no concomitant *MS4A1* mutations could be identified in that sample ^16^.

The propensity of B-cell malignancies to exploit alternative splicing mechanisms of antigen escape is now well-documented, with CD19 and CD22 serving as prime examples ^50,51^. With respect to rituximab treatment, both mutational ^77^ and non-mutational ^20^ mechanism of CD20 loss have been reported, but the former is generally not considered a significant source of rituximab resistance in DLBCL ^78^. Data on post-mosunetuzumab relapses are just starting to emerge ^16^. As more paired pre- and post-relapse samples from patients on CD20-directed therapies become available, the extent to which acquired mutations and aberrant splicing (of both coding and non-coding exons ^79,80^) supplement each other will become clearer.

At least in theory, targeting of aberrant splicing in cancer with small-molecule inhibitors and antisense oligonucleotides (ASO) ^81-84^ is more feasible that reversing hardwired mutations in the DNA. While previous studies have used ASOs to increase translation by blocking inhibitory elements within the 5’-UTR ^85,86^, our results demonstrate that ASOs could also enhance translation by modulating alternative splicing of non-coding exons. Besides CD20, at least a few dozen other genes regulate mRNA translation via alternative splicing of their 5’-UTRs ^87^, and more than 50% of human genes utilize uORFs to regulate protein output in both diseased and healthy tissues ^88^. Harnessing this mechanism to achieve better treatment outcomes will be the next challenge for the cancer immunotherapy community.

## ACKNOWLEDGEMENTS

We are grateful to all members of the ATT laboratory, in particular Priyanka Sehgal, for many helpful discussions. This work was supported by grants from the National Institutes of Health (U01 CA232563 and U01 CA232563-S3 to ATT and YB, U01 CA232486 and U01 CA243072 to SKT, and T32 CA009615 to SZ),

United States Department of Defense (CA180683P1 to SKT), The V Foundation for Cancer Research (T2018-014 to ATT), The Emerson Collective (886246066 to ATT). SKT is a Scholar of the Leukemia & Lymphoma Society. SKT is Joshua Kahan Endowed Chair in Pediatric Leukemia Research and ATT is Mildred L. Roeckle Endowed Chair in Pathology at Children’s Hospital of Philadelphia.

## AUTHORS CONTRIBUTION

ZA, LP, CS, MTD, FX, UZ, YZ, SS, SZ, CDF, JPL, SYY, MA, PKS performed research; ZA, HEK, MML, YB analyzed sequencing data, VP, EC, and SJS analyzed clinical data, SKT, PML, MR and ATT supervised lab experiments; ZA and ATT wrote the paper; everyone else reviewed the manuscript.

