## Supplemental Data and Methods for "Alternative splicing of its 5’-UTR limits CD20 mRNA translation and enables resistance to CD20-directed immunotherapies"

Zhiwei Ang et al

#### Supplemental Data

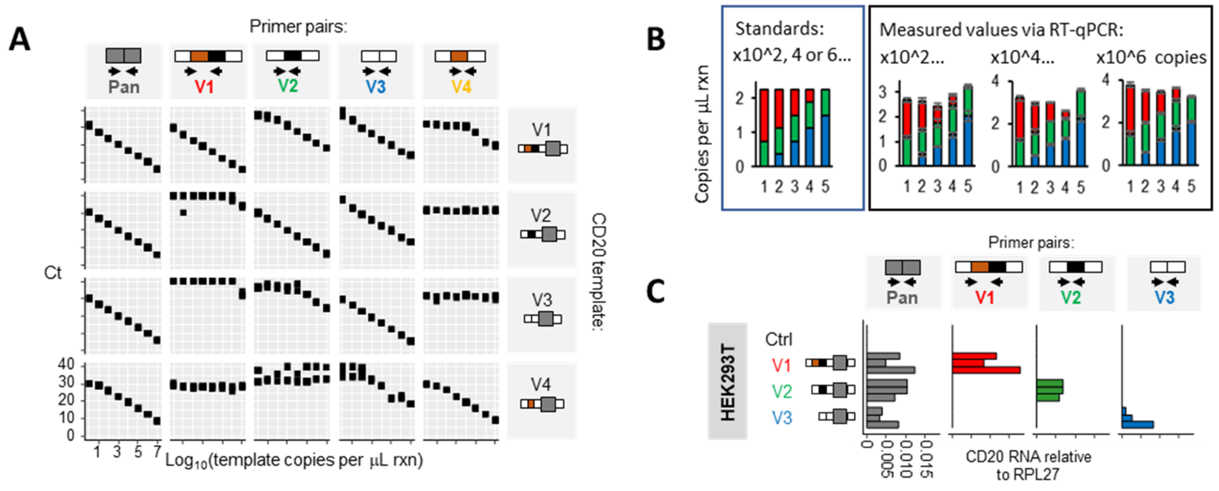


**Supplemental Figure 1. The 5'-UTR-specific primer pairs amplify their intended targets efficiently and specifically. (A, B)** Real-time PCR quantification of single **(A)** and mixed template standards **(B)** using 5'-UTR variant-specific (V1, V2, V3 and V4) and pan-CD20 primer pairs. **(C)** RT-qPCR-mediated quantification of pan-isoform and 5'-UTR variant-specific CD20 mRNAs in HEK293T cells after transient transfection with MXS-CMV-V1, -V2 or -V3 plasmids. In **(B)**, error bars represent the standard deviation of 3 technical replicates. In **(C)**, RNA levels are normalized to the reference gene, RPL27. Each bar or dot is an independent experiment; N=2 for **(A)**; N=3 for **(C)**.

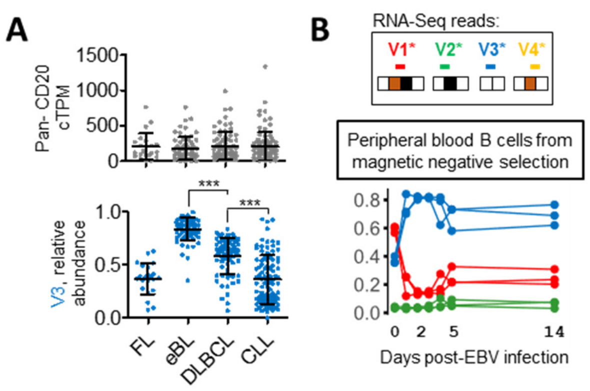


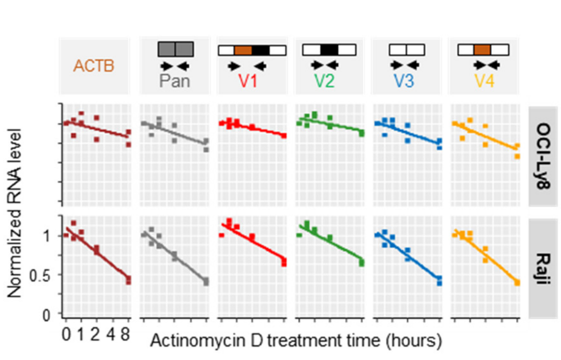


**Supplemental Figure 2. V3 is up-regulated in eBL and in EBV-infected B cells. (A)** Dot plot representing merged values from all datasets for each tumor type displayed in Figure 1H, with mean and SD indicated by whisker plots. *** indicates P<0.0001 per Kruskal-Wallis test (pwc: Dunn test). **(B)** RNA-Seq data of CD20 transcript abundance in human B cells during EBV infection. The relative abundance of V1(red), V2(green), V3(blue) and V4(yellow) is the ratio of sequencing reads mapping to the unique exon-exon junctions found in each 5'-UTR variant of CD20, as color-coded in the "Reads spanning" panel.

**Supplemental Figure 3. The extended 5'-UTRs isoforms, V1 and V2, have longer half-lives.** RT-qPCR-mediated quantification of pan-isoform and 5'-UTR variant-specific CD20 mRNA levels in OCI-Ly8 and Raji cells treated with 10 mg/mL of actinomycin D. RNA levels are normalized to the 0-hour time-point. Each dot is an independent experiment (N=2).

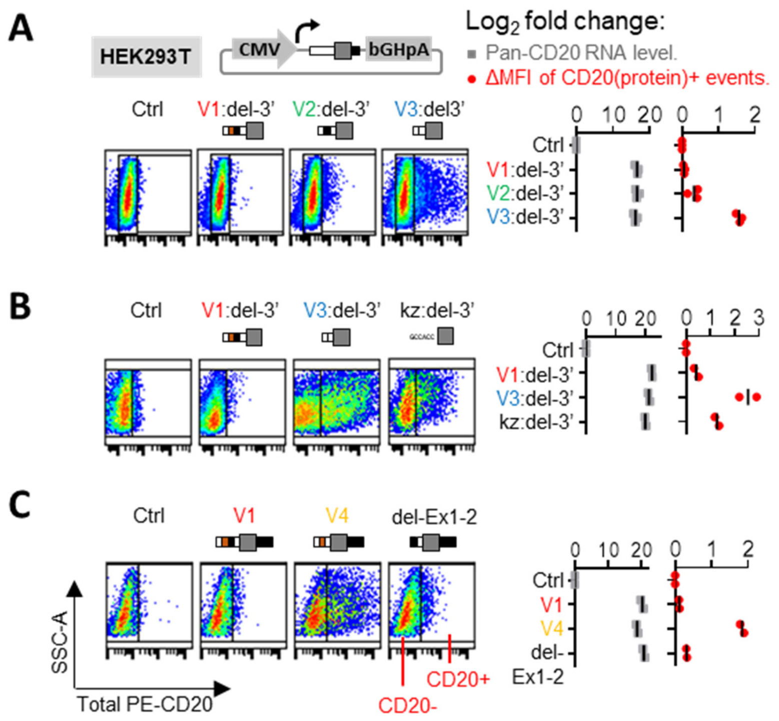


**Supplemental Figure 4. Elements within exon 3a inhibit translation. (A-C)** Measurement of CD20 protein and mRNA levels in HEK293T cells transiently transfected to express CD20 5'-UTR isoforms. Shown at the top right corner of **(A)** is a schematic of the expression vector used. Constructs denoted "del-3'" annotation lack the CD20 3'-UTR sequence. Exons 1 and 2 were deleted from V1 to form del-Ex1-2. Pan-isoform CD20 RNA levels were quantified using RT-qPCR. Total cellular CD20 protein were measured using flow cytometry and expressed as delta median fluorescence intensity (delta MFI = MFI of CD20+ events – MFI of CD20- events). Representative histograms are shown on the left. RNA levels are normalized to the reference gene, RPL27. Each bar or dot represents the average from an independent experiment (N≥2). RNA and MFI values are expressed as log2 fold change relative to Ctrl samples.

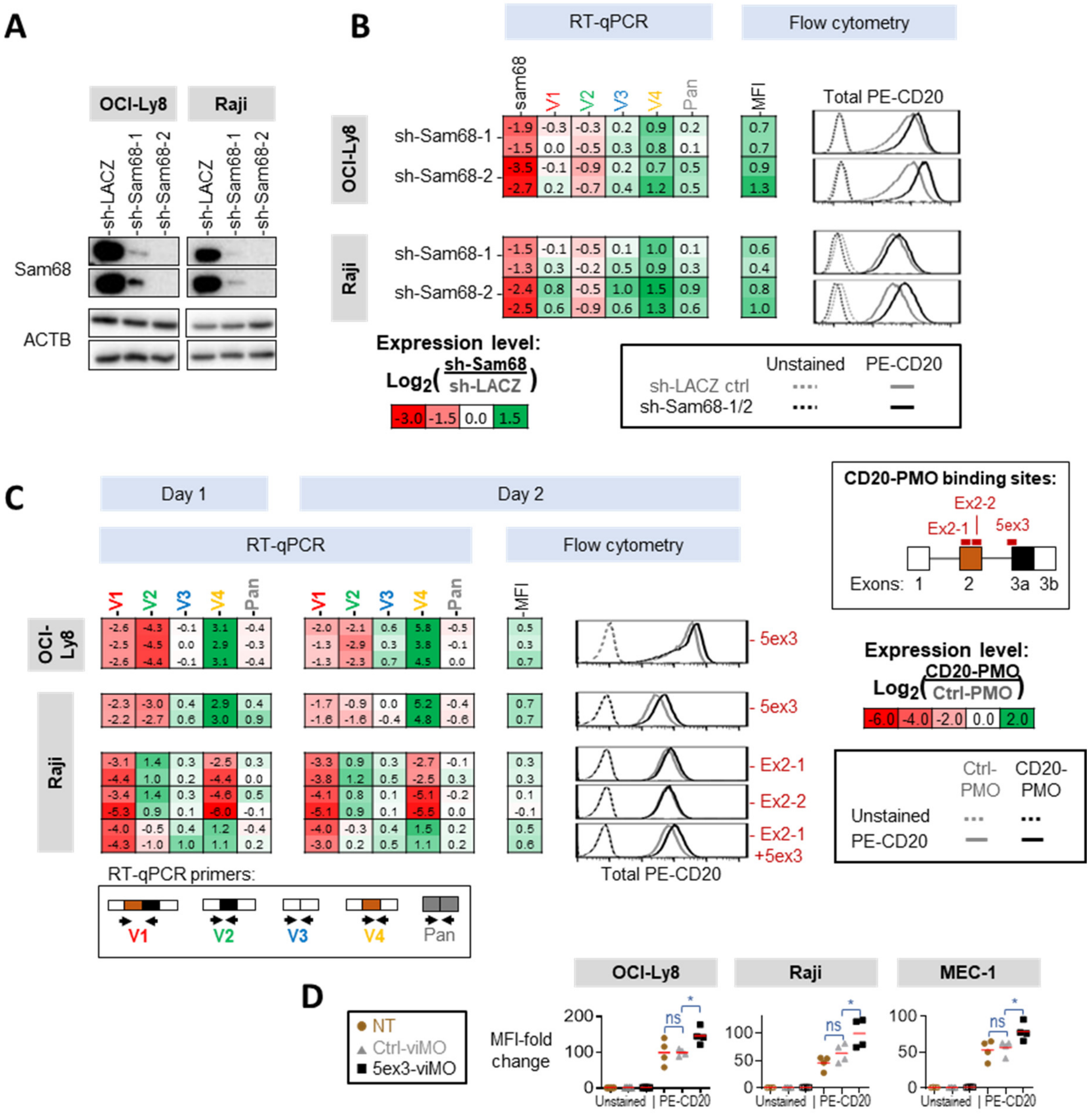


**Supplemental Figure 5. Modulating CD20 splicing enhances its expression. (A-B)** Sam68 and CD20 expression measurements in OCI-Ly8 and Raji cells transduced to stably express either sh-Sam68-1/sh-Sam68-2 or the sh-LACZ negative control. **(C)** CD20 expression measurements in OCI-Ly8 and Raji cells electroporated with various CD20-targeted PMOs: Ex2-1, Ex2-2 and 5ex3. Shown in **(A)** are immunoblots detecting expression of Sam68 and ACTB, which serves as the loading control. Each row is an independent experiment (N≥2). In **(B)** and **(C)**, mRNA levels were quantified with RT-qPCR. The median fluorescence intensities (MFI) of total CD20 protein were determined using flow cytometry. Each heatmap row is an independent experiment (N≥2). Representative flow profiles are shown. **(D)** CD20 protein levels in OCI-Ly8, Raji and MEC-1 cells 2 days after treatment with either the solvent control (NT), Ctrl-viMO, or 5ex3-viMO. The median fluorescence intensities (MFI) of PE-CD20 were determined using flow cytometry and expressed as fold change relative to unstained controls. The * symbol indicates P<0.05 per Kruskal-Wallis test (pwc: Dunn test).

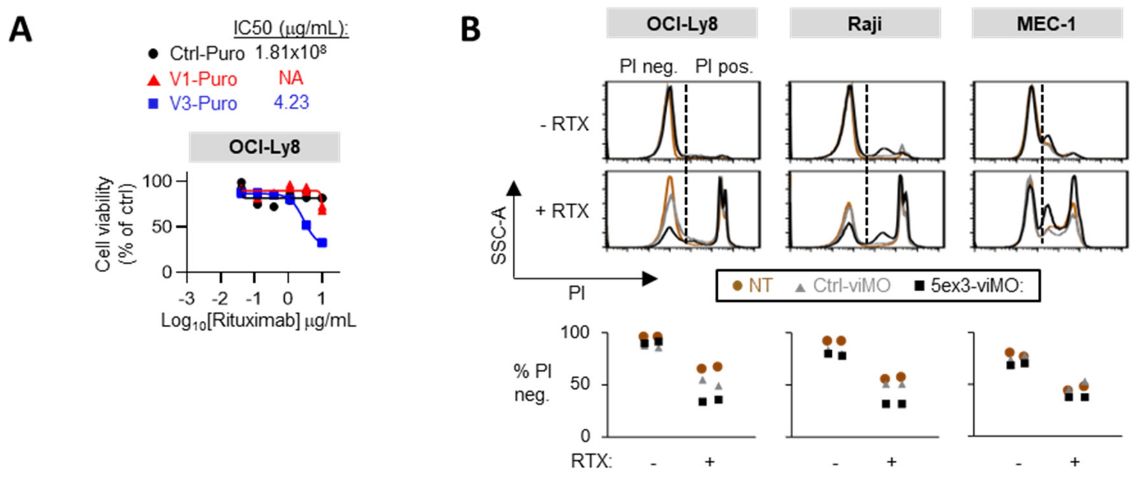


**Supplemental Figure 6. V3-CD20 but not V1-CD20 increases sensitivity to rituximab. (A)** Rituximab dose response curves of OCI-Ly8 cells stably expressing the empty vector (Ctrl-Puro) or the V1 or V3 5'-UTR + CD20 coding sequences (V1- or V3-Puro). Cell viability was measured using the WST-1 assay. All values are normalized to the "no rituximab" control. Each dot is an independent experiment (N≥2). **(B)** Sensitivity to rituximab in OCI-Ly8, Raji, and MEC-1 cells treated with either solvent control (NT), Ctrl-viMO, or 5ex3-viMO followed by 10 μg/mL rituximab. Top: Propidium iodide (PI)-negative, viable cells were detected using flow cytometry. Bottom: Quantitation of data in the top panels. Each dot is an independent experiment (N=2).

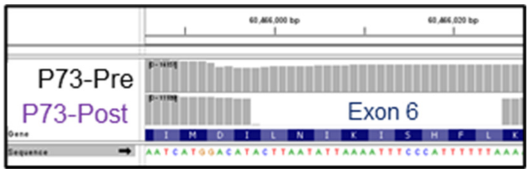


**Supplemental Figure 7: Twenty-eight nucleotide deletion in exon 6 of the *MS4A1* gene in the P73-Post sample.** RNA-Seq read alignments were extracted from and visualized in the IGV viewer.

### Supplemental Methods

**Cell lines and cell culture.** OCI-Ly8 and HEK293T cells were kind gifts from the laboratories of Raju Chaganti (Memorial Sloan Kettering Cancer Center, New York, NY) and Peter Choi (Children's Hospital of Philadelphia, Philadelphia, PA), respectively. MEC-1 cells were obtained from the ATCC (American Type Culture Collection). OCI-Ly8 and Raji cells were authenticated with short tandem repeat profiling. All cell lines were routinely tested to be Mycoplasma free via EZ-PCR Mycoplasma Detection Kit (Sartorius). OCI-Ly8, Raji, and MEC-1 cells were maintained in RPMI-1640 medium supplemented with 10% FBS, 2 mmol/L l-glutamine at 37°C and 5% $CO_2$. The HEK293T cells were maintained in DMEM medium supplemented with 10% FBS, 2 mmol/L l-glutamine at 37°C and 5% $CO_2$.

**Normal and malignant donor B cells.** Cryopreserved normal PBMCs were obtained from the Human Immunology Core, Penn Medicine. Cryopreserved PBMCs from patients with DLBCL, CLL and FL were obtained from the Stem Cell and Xenograft Core, Penn Medicine. These PBMCs were viably thawed and their B cell fractions were enriched using the EasySep™ Human B Cell Enrichment Kit II Without CD43 Depletion (STEMCELL Technologies). After enrichment, more than 90% of the cell populations stained positive for CD19 (via flow cytometry).

**Infection with B95.8 strain EBV.** B95-8 cells (ATCC # CRL 1612; 13) were seeded at 3 x $10^5$ cells/ml in a 75$cm^2$ tissue culture flask in complete RPMI 1640 containing 10% heat-inactivated fetal bovine serum (FBS), Penicillin/Streptomycin at 100U/ml and L-gutaime at 100 µg/ml. After two days, cells were resuspended in fresh complete RPMI 1640 at 1 x $10^6$ cells/ml and stimulated with 20ng/ml tetradecanoyl phorbol acetate (TPA; Sigma-Aldrich, Inc., St. Louis, MO) for 1 hour to induce virus production and then resuspended in complete RPMI for 96 hours before centrifugation at 600 x g for 10 minutes at 4˚C. Cell supernatant was then filtered through a 0.45-micron filter and stored at -80°C. For infection with EBV, 500µl of filtered supernatant containing EBV (B95.8) was added to 2X$10^6$ enriched peripheral blood B cells in 5 ml complete RPMI medium in a 25 $cm^2$ tissue culture flask and placed upright in a $CO_2$ incubator at 37°C for 2-4 hours. Subsequently, 10 ml additional medium and cyclosporin A (1µg/ml; Sigma-Aldrich, Inc., St. Louis, MO) was added. To generate LCLs, the same infection procedure was followed except PBMCs were used instead of enriched B cells. On days three and six, nine, and twelve, half of the medium was replaced with fresh complete RPMI with cyclosporin A. After two weeks, stable EBV+LCLs are formed and anti-CD19 staining was used to confirm that the culture is 100% B cells by flow cytometry.

**FFPE sample processing and RNA-Seq.** In the Phase I/II trial of mosunetuzumab (NCT02500407),[1] patients with relapsed or refractory B-NHL received mosunetuzumab in 3-week cycles, for 8 to 17 cycles depending on tumor response. For our studies, clinical tumor biopsies that had been collected prior to study enrollment (before mosunetuzumab [-Pre]), and after study discontinuation for disease progression (after mostunetuzumab [-Post]) were utilized. By immunohistochemistry (IHC), these FFPE samples were dual-stained with anti-CD20 (clone L26, VENTANA) and anti-PAX5 (clone DAK-PAX5, DAKO) antibodies. Some patients were indicated by pathologist reports to have been CD20 protein positive pre- and negative post-mosunetuzumab. We obtained FFPE samples paired pre and post mosunetuzumab relapse FL tumor samples from four such patients (P29, P73, P100 and P195). RNA was isolated using the Agencourt FormaPure Kit (Beckman Coulter LifeSciences, Indianapolis, IN). Sequencing libraries were prepared using 100ng of RNA and the TruSeq RNA Exome kit (Illumina, Inc., San Diego, CA). Libraries were sequenced on a HiSeq2500 (Illumina) to generate 2x150 bp strand-specific paired-end reads. All samples were sequenced simultaneously, in the same sequencing run. For the cell lines, total cellular RNA was isolated using the Maxwell® RSC simplyRNA Cells Kit (Promega), and 500 ng was submitted to Genewiz LLC, for paired-end RNA-Seq (Illumina, 2x150bp) after Poly(A) selection. All control and treatment samples were sequenced simultaneously, in the same sequencing run.

**Lymphoid tissue processing and RNA-seq.** For the tonsillar B-cells, normal human tonsils from CHOP [2] were manually homogenized with Precision Large Tissue Grinder (Covidien 3500SA) and passed through a 70 µm strainer. Cells were washed with PBS, blocked with Human TruStain

(Biolegend), and stained with anti-CD19 APC (clone HiB19, Biolegend), anti-IgM FITC (Clone G20-127, BD Biosciences), anti-IgD PE (Clone IA6-2, Biolegend). Cells were washed twice in PBS and resuspended in 2% FBS and 0.1 μg/mL DAPI in PBS. CD19+IgM+IgD+, CD19+IgM+, CD19+IgM-, CD19+IgM+IgD-, and CD19+IgM-IgD- cells were sorted on the MoFlo ASTRIOS into 10% FBS RPMI-1640 media with 25 mM HEPES. For BCR stimulations, naïve tonsillar B cells (CD19+IgM+IgD+) from 3 pediatric donors were FACS-sorted and treated with isotype control (Goat F(ab')2 IgG-UNLB, Southern Biotech, Cat.#. 0110-01) or 10ug/mL anti-IgM F(ab')2 (Goat F(ab')2 Anti-Human IgM-UNLB, Southern Biotech, Cat. #. 2022-01) overnight. Viability and proximal BCR signaling were confirmed in cells 20 min after stimulation with various F(ab')2 via phospho-SYK immunoblotting (anti-pSYK, Cell Signaling, Cat. #2710S). From the tonsillar B cell samples, RNA was isolated using Qiagen RNeasy Kit (Micro). RNA integrity and concentration were determined using Eukaryote Total RNA Nano assay on BioAnalyzer. RNA-seq was performed on 10 ng–1μg of total RNA according to the Genewiz Illumina HiSeq protocol for poly(A)-selected samples (2 × 100 or 150 bp, ~350M raw paired-end reads per lane).

**Quantitation of splice isoform.** For this assay, we designed individual PCR primers that spanned unique exon-exon junctions specific for each 5'-UTR isoform of CD20. We also designed a "Pan CD20" primer pair to amplify a common exon-exon junction found in all the known CD20 isoforms. To verify their specificity, we tested these primers against individual as well as admixed PCR products corresponding to all four CD20 isoforms. As expected, the amplification curves were isoform-specific, displaying linear dynamic ranges that spanned eight orders of magnitude with optimal 0.95 to 1.00 PCR efficiencies ($10^{-1/slope} - 1$) for their cognate CD20 templates [**supplemental Figure 1A**]. For example, compared to the cognate V3 template, V1 had to be added in 240-fold excess before the V3-specific primer pair would yield a product at the same cycle threshold (Ct) value. The RT-qPCR assay also accurately measured the relative fractions of each variant within predefined admixtures [**supplemental Figure 1B**] and in HEK293T cells transiently transfected with different CD20 isoforms [**supplemental Figure 1C**].

**RT-qPCR.** Total RNAs were isolated from whole cells using the Maxwell® RSC simplyRNA Cells Kit (Promega). To generate cDNA, 500 ng of RNA (via NanoDrop™ 2000) were hybridized with a mixture of Oligod$(T)_{20}$ and random hexamer primers and reverse transcribed with GoScript™ Reverse Transcriptase (Promega) with a 1-hour extension step at 42 $^{o}$C for OCI-Ly8, Raji and MEC-1 cell line samples; or with SuperScript™ IV VILO™ Master Mix (ThermosFisher Scientific) for all other samples. Quantitative PCR was performed using PowerSYBR Green PCR Master Mix (Life Technologies) on an Applied Biosystems Viia7 machine. According to the Minimum Information for Publication of Quantitative Real-Time PCR Experiments (MIQE) guidelines [3], all primers used had efficiencies exceeding 95% and amplified single PCR products from cDNA mixtures, which was then validated via Sanger sequencing. Reference gene primers used include RPL27 (123 bp amplicon on NM_000988; forward: ATCGCCAAGAGATCAAAGATAA; reverse: TCTGAAGACATCCTTATTGACG), GAPDH (227 bp amplicon on NM_002046; forward: GAAGGTGAAGGTCGGAGTC; reverse: GGAAGATGGTGATGGGATTTC) and ACTB (121 bp amplicon on NM_001101.5; forward: GTCATTCCAAATATGAGATGCGT; reverse: GCTATCACCTCCCCTGTGTG) [4,5]. Other primers used include MS4A1 V1 (147 bp amplicon on NM_152866; forward: TGGAGACTCAGATCCTGAACAA; reverse: CCTCTTCCGAGTGACCTTGG), MS4A1 V2 (64 bp amplicon on NM_152867; forward: AGGCCTTGGAGACTCAGAACTC; reverse: CCTCTTCCGAGTGACCTTGG), MS4A1 V3 (126 bp amplicon on NM_021950; forward: GCCTGGACTACACCACTCAC; reverse: GCTCTCAAAACTCCTGAGTCTCC), MS4A1 V4 (121 bp amplicon on hypothesized V4 of MS4A1; forward: TGGAGACTCAGATCCTGAACAA; reverse: GTCATTTTGCTCTCAAAACTCCTTAT), pan MS4A1 (107 bp amplicon on NM_152866, NM_152867, NM_021950 and V4; forward AAAGAACGTGCTCCAGACCC; reverse: TGTTTCAGTTAGCCCAACCACT) and pan KHDRBS1 (aka Sam68) (95 bp amplicon on NM_006559 and NM_001271878; forward: GGATACCTTTGCCTCCACCT; reverse: GCCTTCGTAGCCTTCGTAACT). Expression levels relative to control were calculated using the 2-ΔΔCT method. When indicated, the data were normalized to the mean CT of GAPDH or ACTB. All other RT-qPCR data were normalized to RPL27. DNA contamination was routinely assessed via qPCR of negative control reverse transcription reactions which lacked reverse transcriptase (cycle thresholds above 35).

**Polysome profiling.** Polysome profiling was performed as described previously [5,6]. Briefly, cells were preincubated with cycloheximide (100 µg/mL, Sigma) for 15 min, and cytoplasmic lysates were prepared and fractionated by ultracentrifugation (at 246000 x g for 3 hours on a SW 41 Ti swinging bucket rotor) through 10%–50% linear sucrose gradients; 18 fractions were collected, and their absorbance at 254 nm (A254) were measured with a NanoDrop™ 2000 Spectrophotometer (Thermo Scientific). Total RNA was isolated in the presence of 4 µg glycogen (Ambion) using TRIzol™ LS Reagent (Ambion). To obtain a DNA-free sample, an additional set of TRIzol reaction tubes were prepared in parallel with no cells added. After phase separation, the upper aqueous phase of the cell-free tubes was removed and replaced with the upper aqueous phase from the cell sample tubes. The new mixture was then subjected to another round of shaking and phase separation to remove any residual DNA. To generate cDNA, the entire RNA pellet was primed with a mixture of Oligod$(T)_{20}$ and random hexamer primers and reverse transcribed with the SuperScript IV First-Strand Synthesis System (Invitrogen) with a 50 min extension step at 55 $^{o}$C. Purified RNAs were stored at -80 $^{o}$C for less than a week before reverse transcription. The cDNA from each fraction was used for quantitative real-time PCR analysis.

**Lentiviral transduction and antibiotic selection.** Lentiviral particles were produced from HEK293T via co-transfection of the transfer plasmids with the psPAX2 and pMD2.G packaging plasmids. The psPAX2 (Addgene plasmid # 12260) and pMD2.G (Addgene plasmid # 12259) plasmids were gifts from Didier Trono. Two days after transduction with these viral particles, Raji and OCI-Ly8 cells were cultured in media containing 1 ug/mL Puromycin or Blasticidin (Invitrogen) for at least two weeks before samples were collected. HEK293T cells were cultured in media containing 10 µg/mL Puromycin.

**Sam68 knockout and knockdown.** Alt-R™ CRISPR-Cas9 sgRNAs (Integrated DNA Technologies or IDT) targeting the TGAACCCTCTCGTGGACGTG and CAAGTACCTGCCCGAACTCA sequences within the Sam68 CDS, were complexed with Alt-R® S.p. Cas9 Nuclease V3 (IDT) and electroporated in OCI-Ly8 cells with the Neon™ Transfection System 10 µL Kit (ThermoFisher) following manufacturer recommendations. Samples were collected between 1 to 3 weeks after electroporation. The pLKO.1-puro lentiviral vectors with predesigned shRNAs which were dubbed sh-Sam68-1 (TRC clone ID: TRCN0000000046; target sequence: GACGGCAGAAATTGAGAAGAT) and sh-Sam68-2 (TRC clone ID: TRCN0000000048; target sequence: GATGAGGAGAATTACTTGGAT), were obtained from Sigma-Aldrich. We stably expressed these plasmids in Raji and OCI-Ly8 cells to knockdown Sam68.

**Flow cytometry.** To measure total cellular CD20 protein, cells were fixed and permeabilized with the BD Cytofix/Cytoperm Fixation/Permeabilization Solution Kit, and blocked with Fc Receptor Blocking Solution (Human TruStain FcX™, Biolegend) prior to staining with PE anti-human CD20 antibodies (clone 2H7, Biolegend). To measure cell-surface CD20, live cells were blocked with Human TruStain FcX™ (Biolegend) prior to staining with APC anti-human CD20 antibodies (clone 2H7, Biolegend). Staining was performed in blocking buffer (PBS with 2% FBS and 1 mM EDTA). Propidium Iodide (PI) was used to exclude dead cells from analysis. The median fluorescence intensity (MFI) of the APC anti-CD20 stained cell populations was normalized to a standard curve generated from Quantum™ APC MESF beads (Bang Laboratories) and expressed as Molecules of Soluble Fluorochrome (MESF). Acquisition was done on a BD Accuri™ C6 Cytometer (BD Biosciences). Manual gating was performed on the FlowJo Software v.10 (BD Biosciences).

**Western blotting.** Total cell lysates were prepared in Laemmli Sample Buffer (Bio-Rad) with 50 mM DTT, resolved on a 4–15% precast polyacrylamide gel (Mini-PROTEAN® TGX™, Bio-Rad) in Tris/glycine/SDS running buffer (Bio-Rad), and then transferred onto a PVDF membrane (Millipore Sigma, Immobilon-P, 0.45 µm) via the Mini Trans-Blot® Cell system (Bio-Rad). These membranes were then probed with antibodies against CD20 (clone E7B7T, Cell Signaling Technology), ACTB (clone 8H10D10, Cell Signaling Technology), and Sam68 (clone 7-1, Santa Cruz Biotechnology, Inc.); followed by anti-rabbit or -mouse horseradish peroxidase–linked secondary antibodies (Cell Signaling Technology). Western blot chemiluminescent signals were captured with an G:Box Chemi XX6 Gel Doc (Syngene).
